# The Virtual Brain Ontology: A Digital Knowledge Framework for Reproducible Brain Network Modeling

**DOI:** 10.1101/2025.11.19.689211

**Authors:** Leon Martin, Konstantin Bülau, Marius Pille, Rico Schmitt, Christoph Hüttl, Jil Meier, Halgurd Taher, Dionysios Perdikis, Michael Schirner, Leon Stefanovski, Petra Ritter

**Affiliations:** Berlin Institute of Health (BIH) at Charité - Universitätsmedizin Berlin, Charitéplatz 1, 10117, Berlin, Germany; Department of Neurology with Experimental Neurology, Charité - Universitätsmedizin Berlin (Corporate member of Freie Universität Berlin and Humboldt Universität zu Berlin), Charitéplatz 1, 10117, Berlin, Germany; Bernstein Focus State Dependencies of Learning and Bernstein Center for Computational Neuroscience, Berlin, Germany; Einstein Center for Neurosciences Berlin, Charitéplatz 1, 10117, Berlin, Germany; Einstein Center Digital Future, Wilhelmstraße 67, 10117, Berlin, Germany

**Keywords:** Keywords: Brain Simulation, Ontology, Knowledge Base, Knowledge Engineering, Semantic Code Generation Framework, Scientific Computing, FAIR research

## Abstract

Computational models of brain network dynamics offer mechanistic insights into brain function and disease, and are utilized for hypothesis generation, data interpretation, and the creation of personalized digital brain twins. However, results remain difficult to reproduce and compare because equations, parameters, networks, and numerical settings are reported inconsistently across the literature, and shared code is often not fully documented, standardized, or executable. We introduce *The Virtual Brain Ontology* (TVB-O), a semantic knowledge base, minimal metadata standard, and Python toolbox that simplifies the description, execution, and sharing of network simulations. TVB-O offers 1) a common vocabulary and ontology for core concepts and axioms representing current domain knowledge for simulating brain network dynamics, 2) a minimal, human- and machine-readable metadata specification for the information needed to reproduce an experiment, 3) a curated database of published models, brain networks, and study configurations, and 4) software that generates executable code for various simulation platforms and programming languages, including The Virtual Brain, Jax, or Julia. FAIR metadata and provenance-aware reports can be exported from TVB-O’s model specification. It hereby enables a flexible framework for adopting new models and enhances reproducibility, comparability, and portability across simulators, while making assumptions explicit and linking models to biomedical knowledge and observation pathways. By reducing technical barriers and standardizing workflows, TVB-O broadens access to computational neuroscience and establishes a foundation for transparent, shareable “digital brain twins” that integrate with clinical pipelines and large-scale data resources.

## 1. INTRODUCTION

Recent advances in computational neuroscience, particularly personalized whole-brain simulations, provide a framework for a mechanistic understanding of complex brain network dynamics in both health and disease, paving the way for personalized treatments [1, 2]. At scale, these simulations generate time series comparable to those of electroencephalography (EEG) and functional magnetic resonance imaging (fMRI) [3]. They are instantiated as networks of mean-field neural population models that capture the average activity within parcellated brain regions [2]. In practice, such personalized models are increasingly used as “digital brain twins” to support hypothesis-driven personalization and clinical reasoning while offering mechanistic mathematical insights into dynamic brain states and their modulation [4, 1, 5, 6]. One of the most widely used simulator software packages for modeling brain activity on a large scale is *The Virtual Brain* (TVB) [7, 8, 9], which produces artificial timeseries data for various modalities, from raw neural activity to simulated neuroimaging data. TVB offers a comprehensive library of neural population models and tools for region-wise and surface-based configurations.

However, adapting or extending this library would require a change in the source code and, hence, does not reflect the need for a flexible way of developing and extending new models. Also, other state-of-the-art whole-brain network modeling platforms, like *neurolib* [10] or *Brain Modeling Toolkit* [11], lack a standardized and user-friendly mechanism and interface for customizing model equations or adding personal models without dealing with the specific programming syntax of these tools. Further, a transfer of model implementations between platforms is not possible, which limits interoperability and reproducibility of results. To enable model extensions or transfer, a high-level, declarative, and interoperable model description language is necessary, one that is both human-readable and machine-actionable. Such a language must also include a clearly defined ontology of modeling entities, guaranteeing clarity and consistency across implementations.

Standardization efforts in systems biology demonstrate the benefits of such approaches. The MIRIAM guidelines (Minimum Information Required In the Annotation of Models) [12] define core principles for annotating computational models with standardized metadata, ensuring clear identification of biological entities and relationships. Similarly, the MIASE guidelines (Minimum Information About a Simulation Experiment) [13] specify the essential information needed to reproduce simulation workflows, including experimental conditions, parameter values, and solver settings. These initiatives have significantly improved the reproducibility and reuse of models in systems biology by providing machine-readable documentation that complements formal model definitions. However, there is no equivalent for whole-brain network modeling. Unlike biochemical networks or cellular pathways, brain network models often depend on nonlinear mean-field dynamics at the population level and incorporate multimodal neuroimaging data. Developing such models requires extensions beyond MIRIAM and MIASE: a specialized, domain-specific language that encodes both the mathematical structure of neural population models and the origin of simulation experiments at the whole-brain level.

When models are published, reproducibility remains limited, partly because of incomplete model descriptions, missing parameter values, ambiguous mathematical formalizations, or unshared or inconsistent code [14, 12, 15, 16]. Merely publishing source code is not enough, since software stacks, dependency versions, and execution environments often differ, causing variability in results [17]. Consequently, a clear metadata framework is necessary to record model definitions, simulation provenance, and experimental context in a standardized and machine-readable way [14].

Although the field of neuroimaging already benefits from strong and established data standards like the Brain Imaging Data Structure (BIDS) [18, 19] and the Neuroimaging Data Model (NIDM) for reporting fMRI analyses [20], current efforts in the domain of computational modelling offer only partial solutions and are rarely adopted in the field.

Emerging efforts each address fragments of reproducible model specification but stop short of a unified semantic and provenance layer. The BIDS extension for computational models [21] standardizes file naming and hashed JSON sidecars for simulation outputs, yet remains a packaging convention without an ontology or formal equation/provenance model. The Low Entropy Model Specification (LEMS) [22] language underpins NeuroML [23], a specification language for biophysically detailed cell and microcircuit descriptions. However, NeuroML and LEMS focus on micro-scale simulations at the neuron level, missing core mathematical elements for brain network modeling and provenance. Data formats like SONATA [24] alleviate performance and size issues but do not introduce higher-level semantic typing. PyRates [25] generates backend-specific code for dynamical analyses (bifurcations, sweeps, fitting) yet lacks an explicit ontology, equation class taxonomy (e.g., stochastic/event-handled systems), and integrated provenance. Consequently, none delivers a semantically explicit, interoperable, and provenance-aware specification for large-scale brain network studies.

As a result, the field lacks a unified, semantically explicit specification for brain network models. This gap leads to implicit assumptions about model components and settings for numerical computation, inconsistent or ambiguous terminology across frameworks, limited opportunities for systematic comparison, and barriers to automated validation against biological data [26]. These issues weaken the FAIR principles of findability, accessibility, interoperability, and reusability [27]. With the rise of large-scale datasets and AI-driven analyses [28, 29, 30, 31], contextualized and programmatic access to models and metadata has become essential. Without established standards, simulation results remain hard to reuse, parameter optimizations cannot be validated across different groups, and community-driven model development slows down. Additionally, simulation outputs are rarely archived systematically, which limits cross-study comparisons and long-term knowledge sharing [32].

Emerging Semantic Web technologies, which provide machine-readable representations of knowledge using shared vocabularies and graph-based data models [33] offer a promising path forward. This approach enables interoperability by linking models and metadata into machine-readable knowledge graphs through a clear semantic description framework, without the need to establish a fully unified terminology. A knowledge graph represents models and experiments as subject-predicate-object triples, connecting them to biological and medical concepts and allowing for reasoning and web-scale interoperability [34, 33]. However, such a semantic framework specifically tailored to the domain of mean-field models and whole-brain network simulations is still lacking.

Therefore, we introduce the Virtual Brain Ontology (TVB-O), a machine-readable semantic knowledge framework for describing mathematical models of brain network dynamics and their solutions and computational derivatives. It combines the mathematics of brain network models with biological and medical contexts and enables automatic generation of executable code and detailed experiment reports. TVB-O offers a high-level declarative language for nonlinear dynamical systems, a semantically annotated library of published models, and tools for creating new models by editing equations or recombining components. Translators produce simulator-specific code and domain-relevant descriptions, while the computational ontology provides an axiomatized vocabulary of all domain-specific entities with their definitions and relations [35, 36]. The TVB-O Python library parses specifications from metadata or natural language to enable dynamic code generation - from refining equations to analyzing dynamical systems - and includes a graphical user interface (GUI) for querying the knowledge graph and exporting executable simulation code with FAIR metadata. Designed for integration of biological processes into mathematical frameworks - from genes and synapses to neurons and whole-brain networks [37, 38] - TVB-O integrates new information into existing knowledge, produces executable models, and supports decision-making in hypothesis-driven neuroscience, thereby advancing FAIR practices through an extensive metadata schema and provenance model.

The model specification framework of TVB-O meets community reporting standards, including MIRIAM and MIASE, and interoperates with related ontologies - such as AtOM [39] for brain atlas metadata, the Gene Ontology (GO) [40] for biological processes, and KiSAO/TEDDY [41] for numerical methods and dynamical behaviors. Being a high-level semantic description of a simulation study, it seamlessly translates to multiple model descriptions and simulation frameworks.

We begin with an overview of the software framework and its machine-readable representation of brain simulation knowledge (Figure 1), then present: (i) a formal ontology that defines the core simulation concepts and relations (Section 2.1); (ii) a minimal metadata scheme for studies and models (Section 2.2); and (iii) a curated database of neural population models and literature-derived simulation experiments aligned to ontology identifiers (Section 2.3). Next, we describe the software stack that supports semantic search in the knowledge base, as well as code generation, execution, and reporting (Section 2.4; Figure 2). Finally, we demonstrate the framework through four use cases that ground the methods in practice: integration with the NeuroMMSig disease knowledge graph, side-by-side comparison of exemplar models, a graph-based BOLD forward model, and an audio-driven stimulus workflow (Section 2.5; Section 2.6; Section 2.7; Section 2.8). We conclude with a brief discussion of implications, limitations, and opportunities for reuse and interoperability.

**Figure 1:**
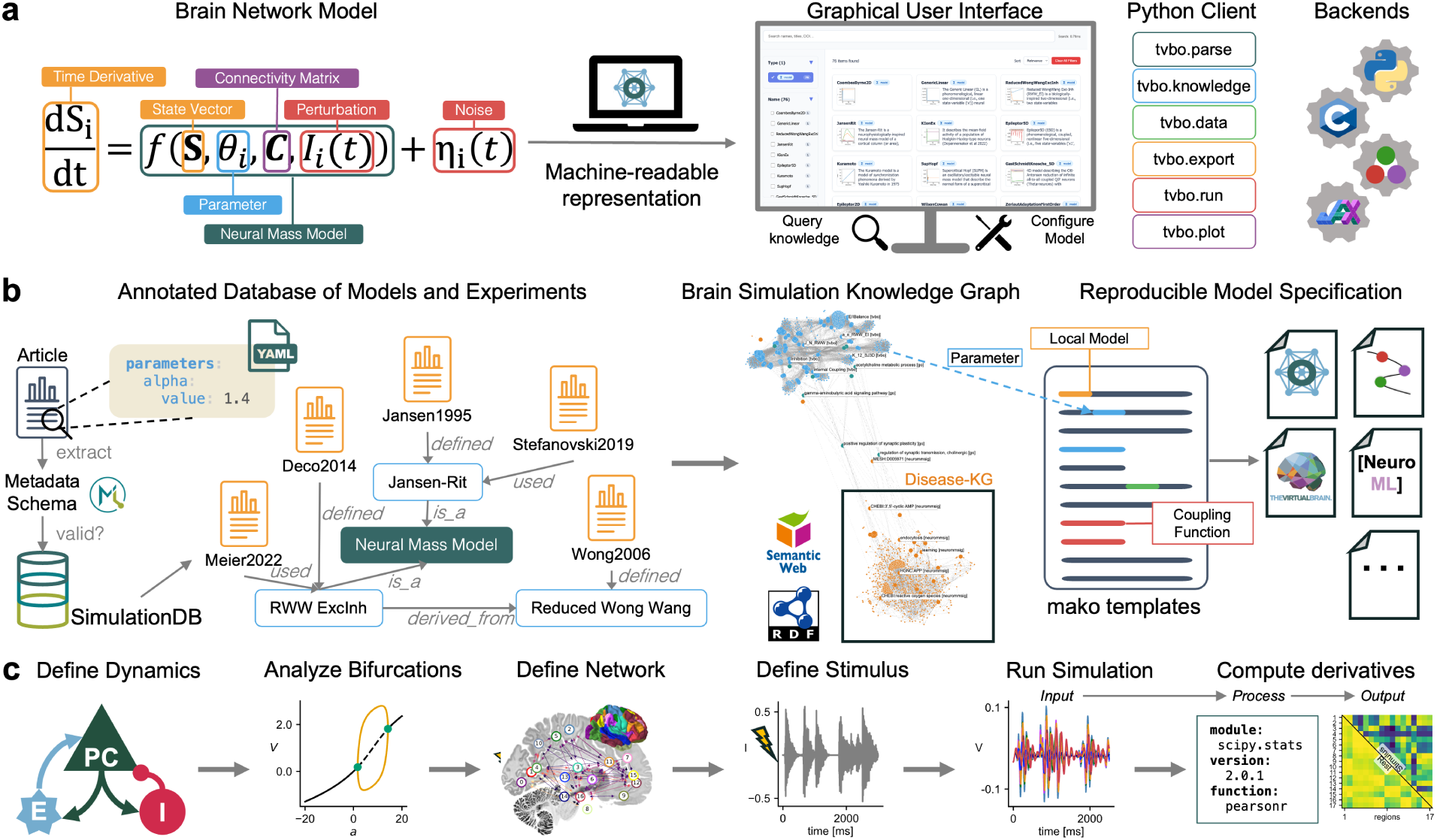
Ontology-driven brain simulation with TVB-O. **a** Our software offers a machine-readable representation of the mathematical framework for brain simulation. The mathematical parts of brain network models are semantically annotated and linked with entities from (non)linear systems dynamics, biology, and medicine. This computational representation of modeling knowledge at a meta-level is language-independent and enables querying, model setup, and interoperability with various programming languages or simulator backends. **b** A curated, annotated database of models and experiments is created by extracting structured metadata from published literature and validating it against an ontology-based metadata schema. These entries are integrated into a knowledge graph that links models to parameters, simulation tools, and disease-related data, ensuring compatibility with the semantic web and supporting reproducible model configuration through templating. Model specifications of the studies shown in the panel are available in the current database; the corresponding citation keys are displayed, and full author lists and publication details are provided in the References [4, 45, 46, 47, 48]. **c** TVB-O guides users through the entire research pipeline of a simulation, including defining local population dynamics, bifurcation analysis, network and stimulus setup, simulation execution, and post-processing of results with forward models or derivatives. Hereby, various simulation platforms and programming languages are supported. Data transformations are shown as directed acyclic graphs of input, process, and output (DAG) to maintain traceability and reproducibility of derived features.

**Figure 2:**
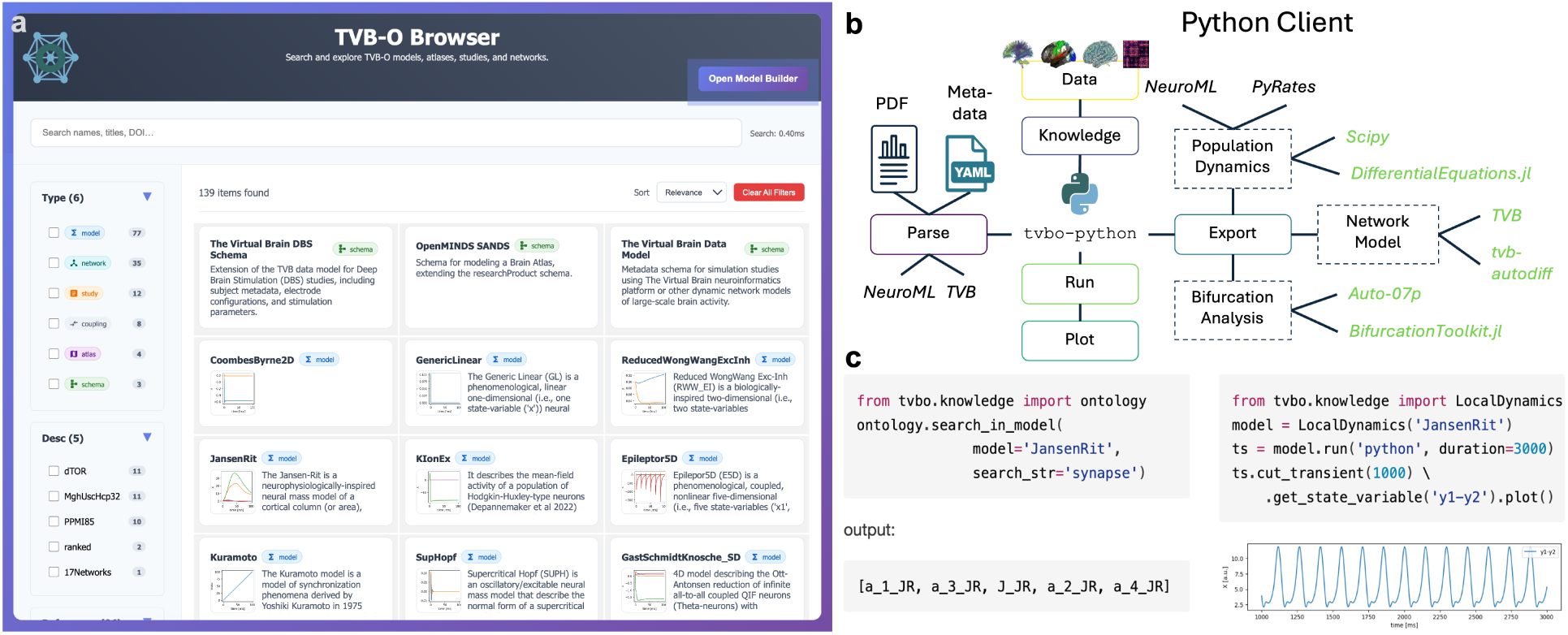
TVB-O Software Framework. **a** The TVB-O Browser graphical user interface (GUI) allows users to search and explore models, atlases, datasets, studies, and brain networks. Models are organized by type, and users can filter by metadata such as descriptions or references. Each entry provides quick access to model details, equations, and reference information, with options to open the model in the builder for further configuration. **b** Detailed overview of software components of TVB-O’s Python client. The core modules include (1) tvbo.parse, loading unstructured text from simulation literature like PDFs, or structured text from metadata files; (2) tvbo.knowledge, integrating information with the knowledge base using axioms defined in the ontology; (3) tvbo.export, converting model specifications into domain-specific simulator code via templating for various software tools (italic); (4) tvbo.run, initiating backends for solving nonlinear systems; and (5) tvbo.plot, offering various visualization functions. **c** Executed code example: the snippet instantiates the Jansen-Rit model from the ontology, performs a semantic search for the term “synapse”, lists the symbols of matching parameters (left), runs a 3 s simulation with the first 1 s discarded as transient, and plots the resulting time series of the derived state variable *y*1 − *y*2 (right), i.e. the net postsynaptic potential difference for the pyramidal population.

In brief, the field faces gaps in specification, semantics, reproducibility, reuse, portability, interoperability, multiscale integration, and automation; TVB-O addresses each with an ontology specification and tools for execution and reuse (see Supplementary Table S1).

## 2. RESULTS

At its core, TVB-O provides a flexible shared vocabulary and metadata schema for describing reproducible whole-brain simulations. Its integration with ontological frameworks of the semantic web allows the use of synonyms and various terminologies while maintaining consistency. This concise language supports custom model equations and research approaches, along with a curated, reusable collection of published configurations and experiments. Such flexibility helps researchers find models and adjustable parameters, examine study data and settings, and reliably reproduce, modify, and share simulations, even when different terminology is used across studies.

The toolkit provides end-to-end capabilities for provenance-backed replication, automated dynamical analyses (including bifurcation), curated inter-regional coupling transforms, surface-based simulations of partial differential equations (PDEs), normative structural connectomes [42, [43], [44]], and cross-backend benchmarking; a consolidated catalogue of curated models and studies is provided in Supplementary Table 1. Technical procedures and architecture are deferred to the Methods and Supplement.

A public web GUI is accessible at https://tvbo.charite.de/ for interactive exploration and configuration. The source code, data model definitions, schemas, and ontology are hosted at https://github.com/virtual-twin/tvbo, with versioned ontology releases available at https://github.com/virtual-twin/tvbo/releases. Docker containers for reproducible environments are available at https://github.com/virtual-twin/tvbo/pkgs/container/tvbo.

### 2.1. TVB Ontology - a minimal, unambiguous vocabulary and logical axioms for brain simulations

TVB-O offers a compact, OWL-based ontology that provides both a controlled vocabulary and formal logical axioms to define and connect the core concepts of whole-brain simulation with consistent identifiers. It includes the essentials-local dynamics (neural mass models with state variables, parameters, and functions), brain networks (parcellations, regions, connectivity weights, and tract lengths), inter-regional coupling (with explicit pre/sum/post semantics), numerical stepping (integrators, methods, delays, noise), inputs (stimuli), and outputs (monitors/observation models and time series) - ensuring that “the same thing” remains unambiguously the same across papers, tools, and code, even when different symbols or synonyms are used. Concepts are structured as classes with specific relationships, like *Connectivity* linked to a *Parcellation*; a *SimulationExperiment* that uses a *Dynamics* model, *Coupling*, *Integrator*, and *Connectivity*, and produces *TimeSeries*. Terms link to persistent IRIs and utilize established vocabularies when suitable: atlas and parcellation entities follow SANDS/OpenMINDS standards [39] and numerical and dynamical concepts are mapped to community identifiers where applicable. This alignment maintains a concise vocabulary while ensuring compatibility with related resources and downstream metadata. This approach makes assumptions explicit when they matter, such as the timing of coupling, which variables a monitor observes, and whether long-range delays depend on tract lengths and speeds. The ontology includes: (i) Neural Mass Models, which describe local dynamics with parameters, state variables, and output transforms; (ii) network connectivity defined by weights and tract lengths tied to a specific parcellation referenced by the AtOM ontology [39]; (iii) coupling functions decomposed into pre-transform, weighted afferent sum, and post-transform, with known formulas; and (iv) an explicit BOLD-like observation model transforming simulated activity. Clarifying these connections allows researchers to pose specific questions (e.g., about hemodynamic observations or sigmoidal coupling) and compare studies despite different notations. For example, [47] refer to a parameter as *J*, which is called *C* in [46]. Such terminology variations are captured within the ontology, enabling flexibility and disambiguation.

The ontology is designed to be minimal by purpose: it establishes a shared language for meaning and relationships without dictating scientific decisions or implementation specifics. More detailed operator forms and mathematical structures are provided in Methods (Section 4.1). The main goal here is to create a common framework that minimizes ambiguity, facilitates comparison across studies, and supports features like search, export, and provenance that are explained below.

TVB-O is integrating anatomy, biology, and numerical methods with established community vocabularies so entries remain comparable beyond this work. Parcellation entities resolve to identifiers in the anatomical UBERON ontology [49], aligning regions across atlases and species and enabling label-agnostic queries over anatomical systems. Model parameters and functions can be annotated with Gene Ontology (GO) [40] processes when their biological role is clear, which supports reproducible traversals from disease knowledge graphs to actionable model handles for hypothesis-driven perturbations. Numerical choices - such as integrator family, deterministic versus stochastic stepping, fixed versus adaptive step size, and tolerances - are mapped to Kinetic Simulation Algorithm Ontology (KiSAO) [50] terms, making solver configurations comparable and portable across different tools. All links use persistent, resolvable IRIs that are stored in the ontology, allowing programmatic reuse and SPARQL queries alongside human-readable reports. This lightweight mapping keeps the TVB-O vocabulary minimal while inheriting community semantics and service resolution from those ontologies. In practice, studies can be filtered by anatomical system (via UBERON), biological process (via GO), or algorithm family (via KiSAO), and exports include both the external identifier and the specific parameterization used in figures, clarifying what changed when swapping backends.

### 2.2. Metadata schema - a compact, ontology-aligned record for reproducible simulations

Along with the ontology, we introduce a minimal, human- and machine-readable metadata schema that captures essential information to determine a simulation outcome. The schema is specified in LinkML (https://github.com/linkml/linkml) and aligned with ontology identifiers, so the same description can be validated, queried, rendered for readers, and translated into simulator-ready code without drift. Concretely, the schema is distributed as a human-readable YAML file and compiled into Python dataclasses for programmatic use; curated examples encoded with it populate the project’s database (cf. Section 2.3).

An experiment record details the local dynamics-listing the model and specifying its parameters, state variables, and optional derived quantities/functions with units and typical domains, with the system type (continuous or discrete) explicitly stated-alongside the brain network, defined by a connectome with node labels, a parcellation tied to a specific atlas and coordinate space, and matrices of weights and tract lengths. It makes inter-regional coupling and delays explicit through equations of pre- and postsynaptic summation processes, deriving delays from tract lengths and conduction speed, and states the numerical method (i.e., integrator, step size, optional stochastic terms, and transients) used to solve the system. Targeted inputs are recorded with their regions, variables, weighting, and timing, while the observation pathway specifies monitors or observation models and any downsampling that maps hidden states to observable signals. Finally, outputs include the resulting time series and, when requested, derived features, each stored along with the settings used to generate them. Each entry binds the essential elements with stable identifiers sufficient to re-instantiate a run and to compare it across tools.

The schema enforces necessary roles, such as requiring a SimulationExperiment to specify a model, connectivity, coupling, integrator, and outputs. It employs concise enumerations like *ImagingModality*, *SystemType*, *BoundaryConditionType*, *DiscretizationMethod*, *OperatorType*, or *ElementType* to keep entries comparable and streamlined. All fields link to persistent identifiers from TVB-O and for atlas resources via AtOM [39], ensuring consistency across different notations. This setup enables precise queries, for example, “experiments stimulating the auditory cortex with a hemodynamic observation,” and guarantees that backend changes do not subtly alter the scientific intent.

The metadata schema benefits both authors and tools: it is easy to read and complete as YAML, yet machine-checkable before ingestion. The same source produces synchronized human-readable summaries, FAIR metadata, and executable configurations. Examples using this schema are distributed with the resource, including models, coupling functions, networks, and study entries in the database directory; the compiled Python dataclasses are stored alongside the codebase for direct programmatic access.

An example metadata instance of a Dynamics class of a Neural Mass Model can be found in the Supplement (Section 1.2.2).

#### 2.2.1. Provenance: linked, validated records make simulation steps traceable and reproducible

TVB-O records provenance as an ontology-backed, W3C PROV-O graph that captures who did what with which resources to produce which outputs (Supplementary Fig. S2). Each run creates a compact directed acyclic graph in which the Simulation Experiment uses the neural mass model and its parameters, the brain network and coupling, the integrator and method, initial conditions, and any stimulus, generating raw time series (prov:used, prov:wasGeneratedBy). A downstream monitoring function or forward-model activity consumes the raw series and produces physiological signals; observation/derivative steps are linked to those outputs from which they are derived (prov:wasDerivedFrom). Local and global connectivity are recorded as separate entities, with their aggregation being explicit; their preparation steps (surface reconstruction, tractography, parcellation) are attributed to external software and cited references. Parameter and state-variable derivations are documented so that reported values clearly correspond to the implemented quantities. All nodes are labeled with stable identifiers from the TVB-O ontology (Entity, Activity, Agent) and validated against the metadata schema that enforces roles and domain-range constraints. The result is FAIR, machine-actionable provenance that supports precise queries, re-execution from a single specification, and verifiable figures without relying on narrative descriptions. TVB-O imports PROV-O and reuses its core classes and properties, specializing domain concepts as subclasses that inherit PROV semantics (for example, *SimulationExperiment* ⊑ *prov:Activity*; *NeuralMassModel*, *Connectivity*, *TimeSeries* ⊑ *prov:Entity*; Software and Author ⊑ *prov:Agent*). We use PROV relations directly (*prov:used*, *prov:wasGeneratedBy*, *prov:wasDerivedFrom*, *prov:wasAttributedTo*); when role specificity is needed, we attach roles via qualified associations (*prov:qualifiedUsage* with *prov:role*) to distinguish, for instance, a coupling function from an integrator within the same activity. All terms and instances resolve to persistent IRIs in the TVB-O namespace. Schema-level constraints check domains, ranges, and cardinalities. Provenance thus lives inside the same knowledge graph as models and metadata, enabling seamless SPARQL queries and export to machine-readable serializations, like JSON-LD [51] alongside human-readable reports from a single source.

### 2.3. Model database - curated, versioned resources for retrieval, comparison, and reuse

TVB-O distributes a curated, versioned collection of literature-based study specifications and canonical model definitions. Currently, the database comprises 12 study entries and 78 local-dynamics definitions (24 canonical neural mass models 54 additional continuous and discrete nonlinear Systems), together with 8 reusable coupling functions, 4 parcellated atlases, and 35 structural networks from 19 to 1000 Nodes. Coverage spans widely used neural-mass families (e.g., Jansen-Rit; Breakspear/Larter-Breakspear; Wong-Wang variants; Tsodyks-Markram; Hopfield; Wilson-Cowan; Kuramoto; Epileptor; Montbrió-Pazó-Roxin; generic oscillators) and includes both resting-state and task-style experiments. A comprehensive table with provenance details and per-model characteristics is available in Supplementary Table 1.

Normative structural connectivity in the database comes from three normative tractogram aggregates: (i) 32 adult subjects [52, 53]; (ii) 85 PPMI Parkinson’s disease patients [54, 44]; (iii) 985 HCP young adults (dTOR-985) [42]. Each is parcellated across standard template-space atlas parcellations of increasing granularity (Overview in Supplementary Table 3), producing symmetric streamline count (weight) matrices and mean tract length matrices (mm) suitable for delay derivation. All atlas-connectome pairs are minimally provenance-tagged, internally validated (labels, shapes, units), and versioned via TemplateFlow resources [55]. They provide interchangeable, ready-to-use starting networks for reproducible whole-brain simulations, where original connectomic study data are missing.

Each entry links the published model description to its key parameters, the structural network used, any reported stimuli, and the recorded outputs. This allows users to: (i) retrieve parameter settings as they appeared in the source; (ii) compare how different studies configured the same underlying model; and (iii) rerun or modify an experiment with a clear understanding of what would change. New entries are versioned rather than overwritten, so earlier configurations remain citable. As the community adds more models (e.g., additional Epileptor variants or biophysical observation pipelines), the collection can expand without disrupting existing references.

In practice, researchers can treat the database as a starting shelf: select a model-network pair, examine its adjustable parameters, and either reproduce the published run or initiate a modified study with different inputs or outputs. The underlying symbolic forms support later analytical tasks (such as exploring dynamical regimes), but readers only need the high-level summary to start reuse.

### 2.4. Software - GUI and Python client for search, code generation, and execution

#### 2.4.1. Graphical User Interface: TVB-O Browser and Model Builder

We present an interactive GUI that brings together mathematical models of dynamical systems, simulation studies, brain parcellations, structural connectivity, and coupling functions from the TVB-O knowledge base into a single, coherent browser. The interface and its growing database are available as a web application at https://tvbo.charite.de, providing broad access for the community. By consolidating previously disparate resources that span multiple model families and application contexts, it enables rapid discovery, comparison, and triage across domains that are commonly explored in isolation.

In a unified search view, users can query names, concepts, and citation information, then refine results with context-aware filters automatically tailored to the available content. Each resource type is summarized with details that are immediately useful for scientific assessment. For dynamical systems models, summaries cover the number and identity of parameters and state variables, the presence of interactions or coupling terms, and, when requested, concise narrative overviews with small, standardized preview traces. Brain parcellations and structural networks are characterized by regional granularity and accompanied by compact visual previews that convey structure at a glance. Simulation studies surface citation details to support attribution and literature navigation. Together, these features lower the barrier for new users while allowing experts to scan and compare alternatives quickly and consistently.

An integrated Model Builder extends the browser from passive exploration to active composition. Within a guided dialog, users select a base dynamical system, adjust parameters, define state equations and derived quantities, specify coupling and output transforms, and choose among empirical or synthetic networks where relevant. Inline mathematical previews support verification of expressions prior to export. The builder produces a structured model description together with concise, executable starter material, enabling rapid prototyping, teaching, and reproducible sharing of model configurations assembled in the browser.

Users either pick a published model from the library or add their own. They then choose a brain network, optionally add an input (e.g., a stimulus), and decide what signals to record. Each choice is checked for consistency and can be exported or reloaded later; there is no need to rewrite formulas or glue code.

Together, these components operationalize ontology-based model specification into a usable workflow (Figure 2). The GUI surfaces ontology and study concepts through semantic search and constrains selections to valid combinations, reducing configuration errors. For example, the Jansen-Rit model [46] requires a sigmoidal coupling function to prevent numerical overflow. Each workspace selection materializes into a schema-validated specification that can be exported as runnable code, FAIR metadata, and a human-readable report from the same source. In practice, using a language-independent formal model specification coupled with automated code generation ensures that all modelling decisions remain explicitly traceable from curation to execution. Further architectural details are provided in Methods (see Section 4.4.1 to Section 4.4.9).

### 2.4.2. Python client: templating, execution, and reports

#### 2.4.3. One description produces synchronized human-, machine-, and simulator-ready outputs

The same concise description behind a study can produce human-readable summaries, machine-readable records, and runnable configurations for different simulation tools. This “single source” prevents the familiar drift between a paper’s methods text, a separate code repository, and a Supplementary spreadsheet.

#### 2.4.4. Built-in reports render equations and parameters from the same specification

Figure 3 illustrates this flow: a minimal text description becomes (i) a readable model sheet and (ii) ready-to-run components. Because all outputs are derived from the same source, updates propagate automatically and inconsistencies are avoided.

**Figure 3:**
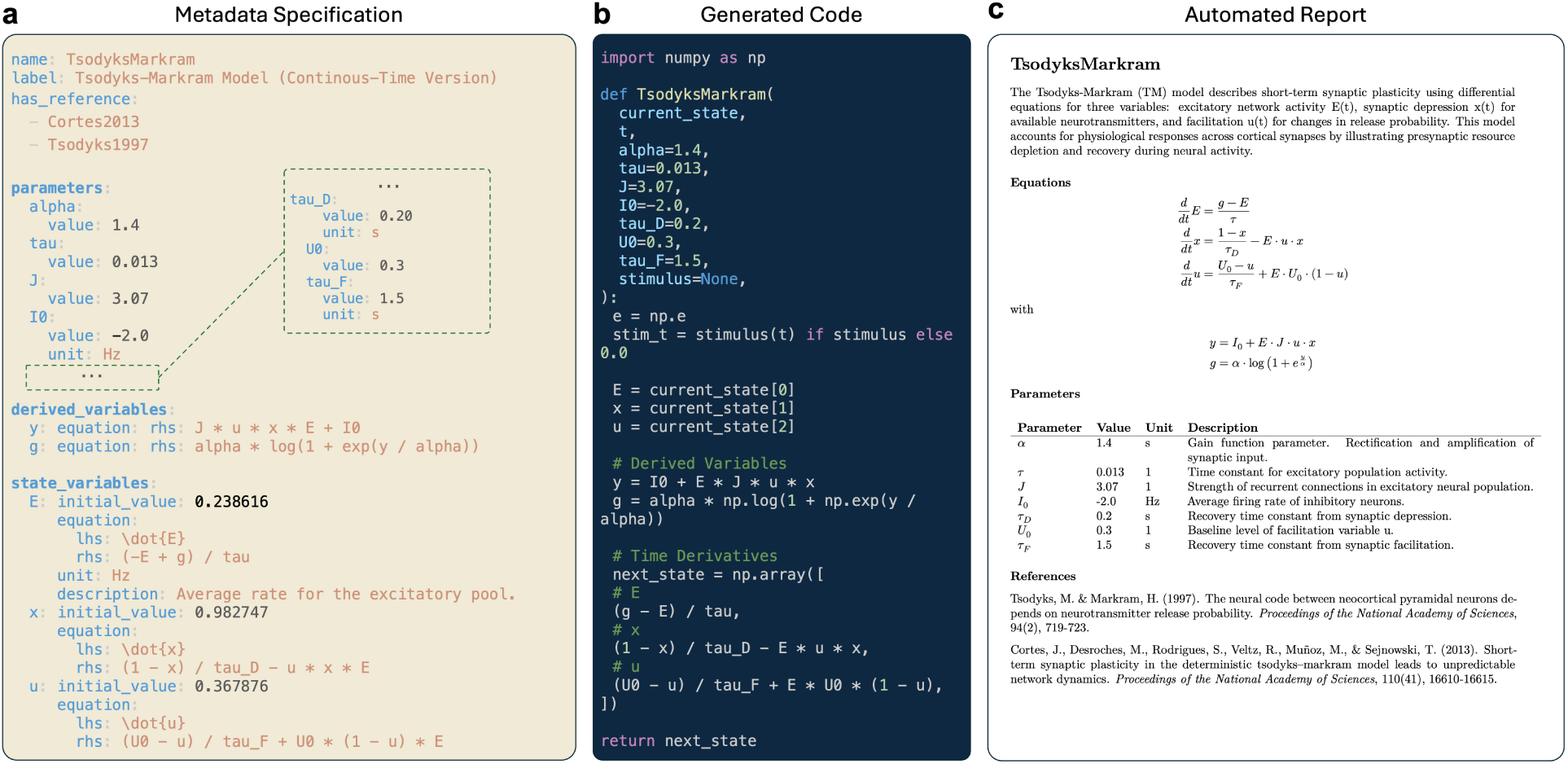
Model specification and export pipeline. **a** Minimal metadata file describing the continuous-time version of the Tsodyks-Markram model [56]. A YAML file with key-value pairs specifies parameters, derived variables, and state equations. **b** Attributes and expressions are parsed and integrated into the ontology and can be exported as a Python function of the dynamical system. **c** Due to the symbolic representation of the model’s mathematics, equations used for creating the code can be rendered in html, LaTeX or markdown, to generate a model report with parameters and references.

#### 2.4.5. Interoperability

TVB-O enables interoperability with the TVB Simulator by linking each simulation component - including local dynamics, coupling, network, integrator, stimulation, and monitors - to stable ontology identifiers. A configured TVB Simulator can be seamlessly integrated into a semantic SimulationExperiment; the exact specifications can then be exported as (i) simulator-ready code for various numerical backends (such as differentiable JAX kernels, PDE surface solvers, LEMS/RateML artifacts), (ii) FAIR metadata, and (iii) human-readable reports. Backend templates only operate on ontology-linked symbols (such as state variables, parameters, delays, and observation pathways), ensuring semantic consistency when changing the implementation language, solver type, numerical precision, or hardware acceleration.

This reversible translation preserves the scientific intent while enabling gradient-based exploration, parameter inference, cross-platform benchmarking, and controlled perturbation studies without re-implementing model logic. Modified configurations can be transferred to TVB or other environments with provenance maintained, supporting side-by-side evaluation of numerical assumptions and biologically grounded parameter adjustments across platforms.

### 2.5. Use case 1: TVB-O enables reproducible, automated routes from disease knowledge to model parameters

We anchored TVB-O modeling entities to the Gene Ontology (GO) to enable reproducible traversals from biology to model parameters. In the current release, the ontology defines 441 explicit links from 210 TVB-O classes to 217 distinct GO terms via OWL restrictions. Most of these (394) use the *surrogate_of* predicate, connecting parameter and state level constructs to biological processes (e.g., Parameter *A* of the Jansen-Rit Model [46] to excitatory postsynaptic potential and *B* of the same model to inhibitory postsynaptic potential, or *C* of the Cakan-Obermeyer model [57] to GO:“stabilization and regulation of membrane potential”). Higher-level regional constructs add 47 extra links via *regional_surrogate_of* (e.g., E/I Balance and Feedback Scaling to acetylcholine metabolism and neuropeptide-mediated retrograde signaling), offering conceptual anchors for regional dynamics. Coverage includes ModelParametersCatalogue, concrete Parameters, and StateVariables, creating an extensive, machine-queryable connection from model structure to curated biological functions. All models are linked to their original publications introducing the dynamics.

For a first showcase of the interoperability of TVB-O with other Knowledge graphs, we connected to TVB-O the disease-focused knowledge base NeuroMMSig [58] - a mechanism-enrichment resource that encodes essential pathophysiology mechanisms of neurodegenerative diseases and supports mechanistic interpretation of multiscale, multimodal clinical data via shared GO processes, enabling systematic hypothesis generation (Figure 4). Starting from a pathology term or disease module, shortest paths through GO to TVB-O identify biologically grounded “handles” in the model (e.g., processes affecting excitation-inhibition balance) that can be perturbed in silico. Because all nodes resolve to stable identifiers, these traversals are reproducible queries rather than ad-hoc mappings, ensuring consistent transfer of evidence from disease biology into model setup.

**Figure 4:**
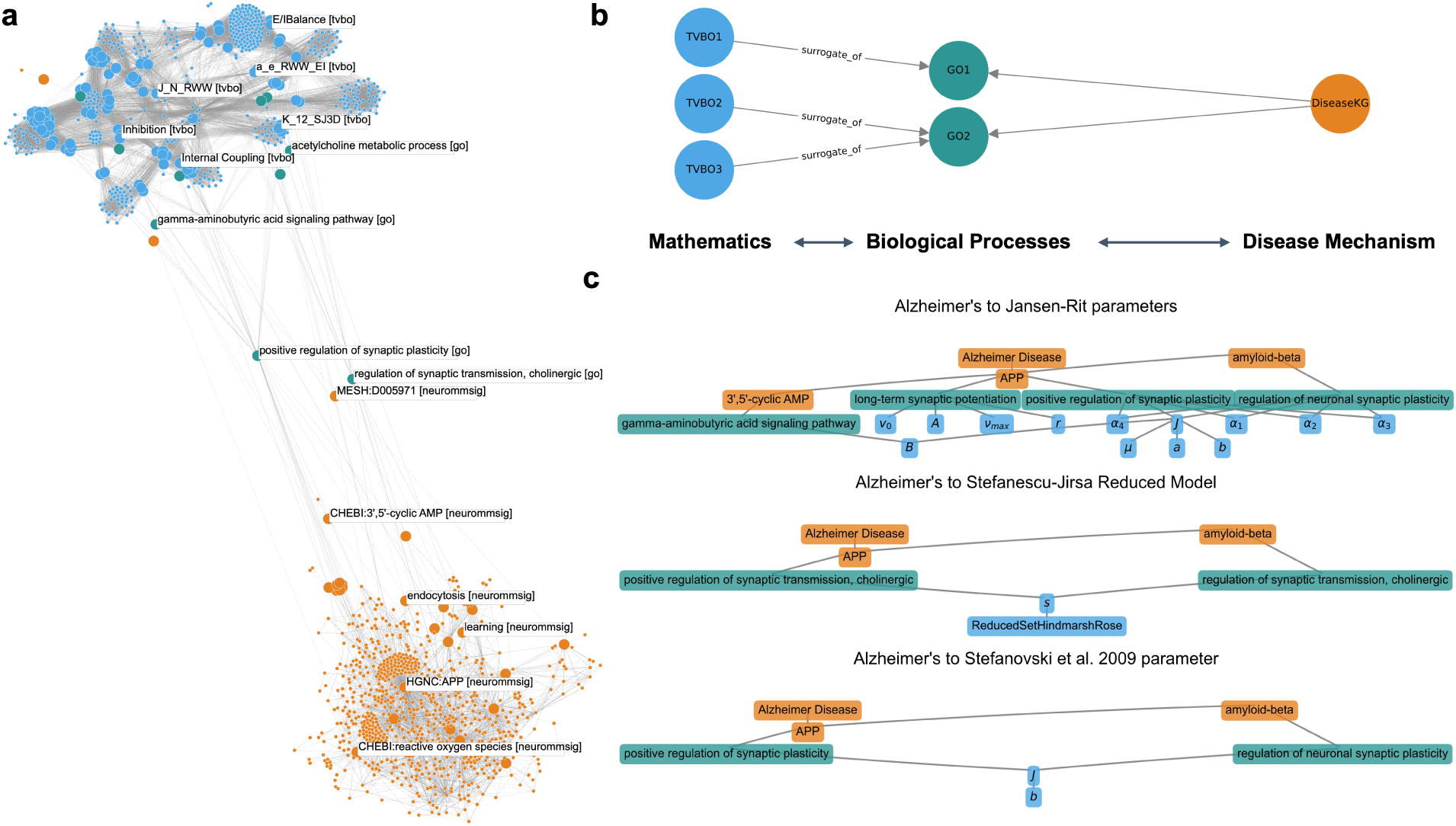
Semantic Interoperability with Disease-Specific Knowledge Graph. **a** The connections between TVB-O (blue) and NeuroMMSig, a disease-specific knowledge graph for neurological diseases (orange), are shown by nodes that directly link the graphs, which are enlarged for emphasis. Both graphs are subsets of the full networks, containing only direct neighbors (small nodes). The bridges are formed by biological processes described by the Gene Ontology (GO) that are present in both graphs. **b** Illustration of bridging mathematics with pathology through biological processes. **c** Example of the shortest pathways from Alzheimer’s Disease in NeuroMMSig to mathematical parameters in TVB-O.

**Figure 5:**
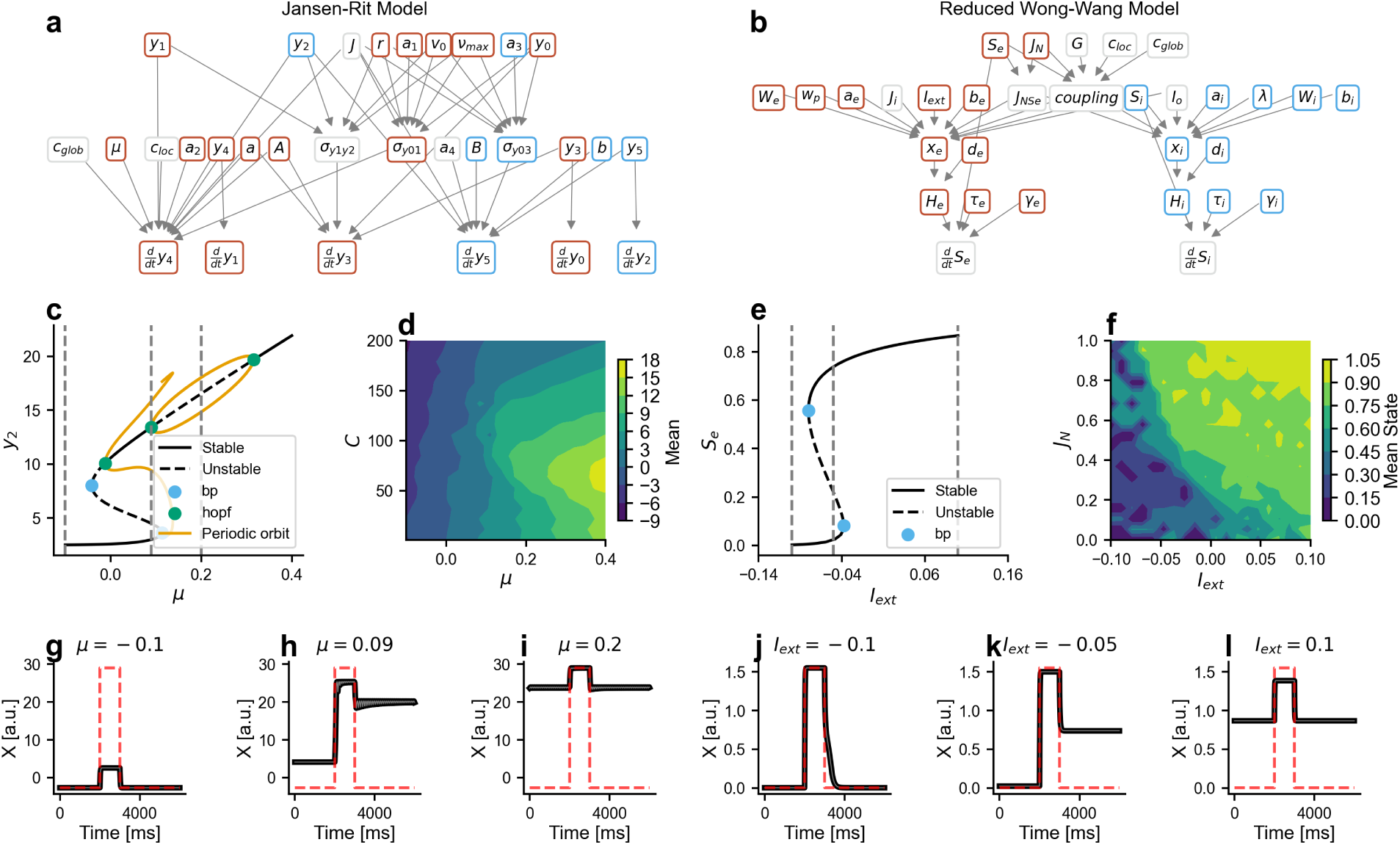
Comparing excitability parameters in two neural mass models with TVB-O. **a-b** Dependency graphs reveal parameter-state variable interactions for Jansen-Rit and reduced Wong-Wang models, with excitatory (red) and inhibitory (blue) nodes. **c,e** Bifurcation diagrams show stable fixed points (solid), unstable equilibria (dashed), and periodic orbits (orange), revealing three regimes: non-excitable, bistable, and overexcited states. **d,f** Contour maps demonstrate how excitability and stimulus amplitude jointly determine mean output across parameter space. **g-l** Time series illustrate regime-specific responses to pulse-train stimulation, with stimulus timing overlaid (red dashed).

### 2.6. Use case 2: TVB-O reveals shared dynamical regimes across diverse models

To demonstrate how TVB-O facilitates cross-model comparison, we analyzed excitability-dependent dynamics in two structurally different neural mass models: the Jansen-Rit [46] and reduced Wong-Wang [45, [48]] models. Both models were instantiated from the ontology, with dependency graphs automatically generated to visualize parameter-state variable interactions colored by excitatory/inhibitory character, as queried from the parameter descriptions. Bifurcation analysis via automated continuation revealed three dynamical regimes as functions of excitability parameters (*µ* for Jansen-Rit, *I*_ext_ for reduced Wong-Wang): 1) non-excitable fixed points at low values, 2) bistability with coexisting quiescent and 3) active states at intermediate values, and overexcited dynamics (saturated activity or limit cycles) at high values. Contour analysis across 8s simulations (2s transient discarded) mapped mean output as joint functions of excitability and stimulus amplitude (*µ* vs. *C* for Jansen-Rit; *I*_ext_ vs. *J_N_* for reduced Wong-Wang), revealing how intrinsic excitability and external drive combine to shape steady-state responses. Time series from single-node simulations at three representative parameter values, with single pulse stimulation (onset 4 s, duration 1 s, amplitude 0.05) applied to designated variables (*y*_4_ for Jansen-Rit, *S_e_* for reduced Wong-Wang), confirmed transitions between regimes by external perturbation. All analyses-bifurcation diagrams, parameter sweeps, and time series-were generated from the same symbolic specifications using TVB-O’s unified export and execution pipeline.

Despite using different mathematical forms, both models show similar shifts in behavior as a key “excitability” setting is increased (Supplementary Figure 1). Comparing their regimes side by side helps readers understand that similar control knobs exist, even when notation varies, and emphasizes where external input can trigger switches between quiet and active states.

### 2.7. Use case 3: TVB-O standardizes observation pipelines for transparent, reproducible transformations

Every step that turns raw simulated activity into something we interpret (like a BOLD-like signal or connectivity matrix) is recorded and repeatable. Making these transformations explicit (see Figure 6) elucidates the processes occurring between the “model output” and the “figure in a paper”. Readers are able to discern (and replicate) how neural activity was transformed into a BOLD-like signal, as well as how summary measures were derived. The distinctions between raw and transformed connectivity emphasize that observational choices impact the conclusions drawn.

**Figure 6:**
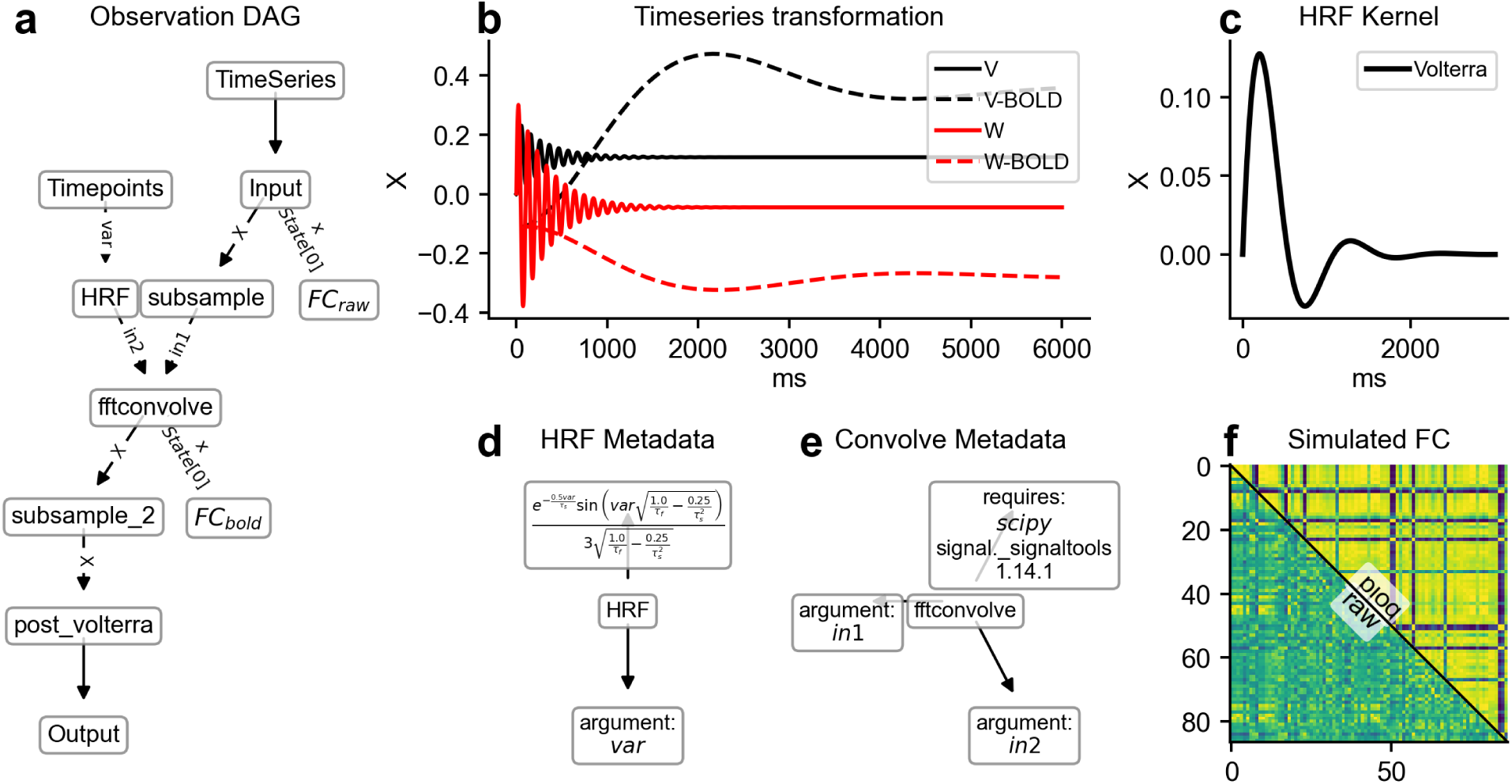
Flexible and modular observation models and their derivatives. Observation models convert simulated neural activity into observable signals through reproducible computational graphs. (a) A directed acyclic graph (DAG) representation of a BOLD forward model uses modular components (nodes) to transform neural time series into BOLD signals as measured by fMRI. (b) The time series transformation is illustrated by convolving simulated neural signals V and W with the hemodynamic response function (HRF) to generate corresponding BOLD signals. (c) The Volterra kernel is used as the HRF. (d) Metadata for the HRF node specify the time-series data of the state variable as the input argument. (e) Metadata for the convolution node describe the implementation with scipy.signal.fftconvolve, which uses Fast Fourier Transform convolution. (f) The output of the derivative node FCbold shows functional connectivity (FC) computed from raw (lower triangle) and BOLD-convolved (upper triangle) signals. Derivatives are endpoints of the DAG and therefore result in a different dimensionality compared to the original time series data.

**Figure 7:**
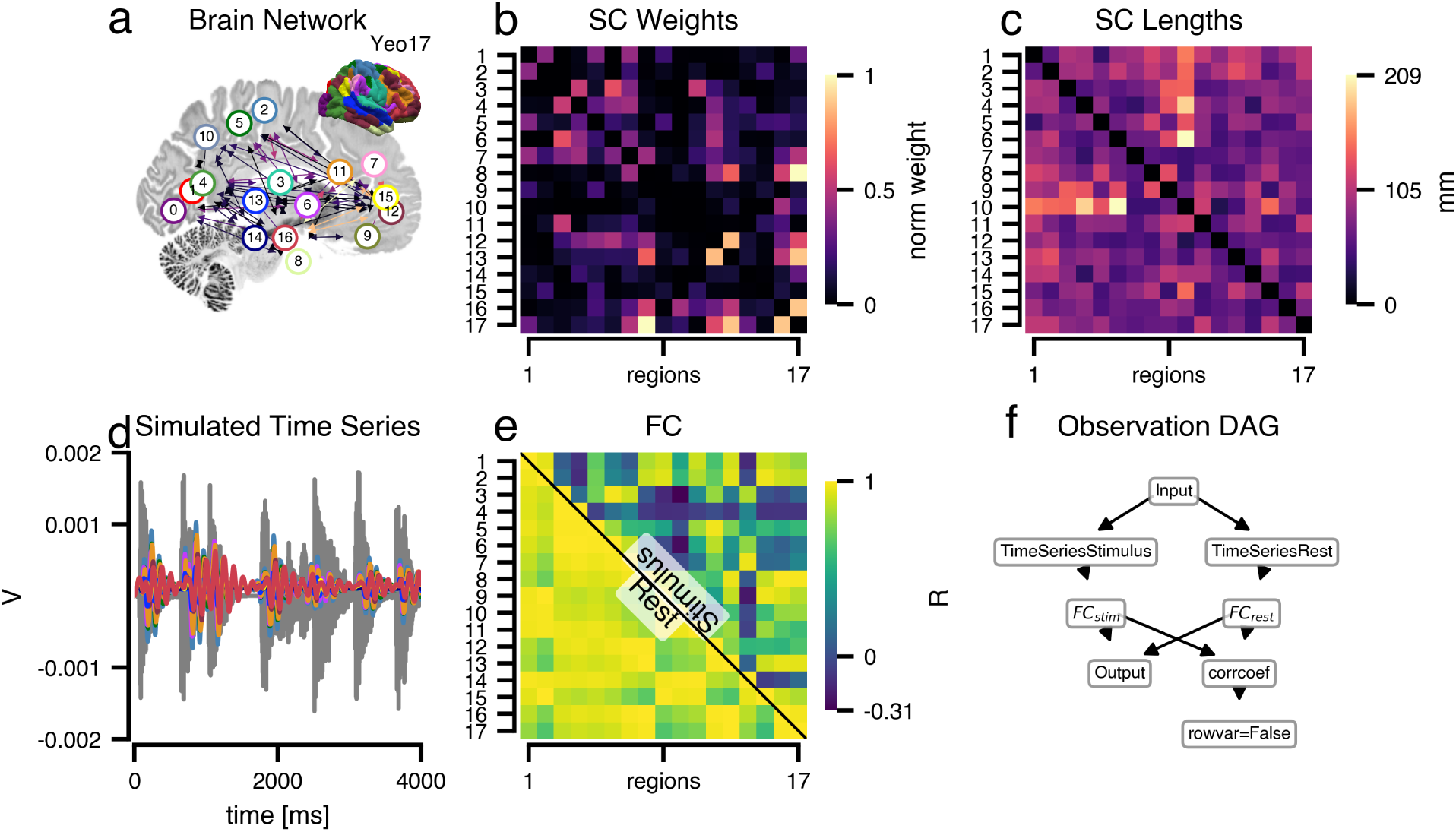
TVB-O enables complex stimulation patterns. Acoustic stimulation of a single node and its network response was simulated using coupled generic 2D oscillators. **a** Structural brain network based on 17 functionally distinct areas from the Yeo parcellation overlaid on an anatomical slice with an inset cortical surface showing the functional networks. An acoustic stimulus was applied to node 3, representing the ventral somatomotor network where the primary auditory cortex is located. **b+c** Heatmaps of normalized structural connectivity weights and tract lengths derived from TVB-O data, indicating the amount of streamlines connecting two regions and the average lengths of those streamlines. **d** Simulated regional time series for nodes connected to the stimulated node (node 3), with the scaled stimulus waveform overlaid in gray for temporal reference. **e** Composite functional connectivity matrix of the simulation without stimulus (lower triangle) and with stimulus (upper triangle). The observation model directed acyclic graph illustrates the dual-input processing pipeline from raw time series to functional connectivity derivatives. Rendering used the project’s custom plotting theme (bsplot with tvbo style); simulations returned labeled time series with dimensions (*time × state × region × mode*) and were generated from the same symbolic specification that underlies code export, bifurcation analysis, and human-readable reporting. **f** Provenance graph indicating the dataflow of the observation models computed in this experiment.

### 2.8. Use case 4: TVB-O makes network perturbations reproducible via formally specified stimuli

The updated metadata framework allows for the inclusion of more complex stimuli and enables precise specification of the model variables that these stimuli influence. Previously, such flexibility was not possible due to inherent limitations in the original implementation. These modifications not only address these constraints but also improve experimental control and increase the model’s versatility.

In this use-case, we injected a complex acoustic stimulus into the ventral somatomotor network, including the primary auditory area [59]. The response propagates along known anatomical connections, and the pattern of inter-regional relationships shifts compared to rest. Since the stimulus is formally recorded (including its location and waveform), anyone can replicate or modify the scenario without reverse-engineering hidden code.

### 2.9. Optimization workflows - ontology-backed extensions of curated experiments

TVB-O’s export pipeline feeds directly into the TVB-Optim parameter optimization suite [60], allowing any curated *SimulationExperiment* to become a differentiable simulator while preserving ontology-backed semantics. Requesting the JAX pathway produces a compiled kernel together with a structured state that includes the initial conditions, connectome, parameterization, stimuli, and observation models exactly as curated in the knowledge base. Because no hand-written glue code is required, optimization experiments inherit the same identifiers, units, and provenance metadata that governed the original curation.

TVB-Optim wraps this state in JAX-native parameter objects and exploration axes that support systematic searches across scientifically meaningful ranges. Grid, uniform, or distributional axes can be assigned to any ontology-referenced quantity, and the resulting spaces are automatically reshaped for vectorized execution across CPUs, GPUs, or multi-device backends. Sequential and parallel execution engines then evaluate these combinations or launch batched runs via accelerator-aware primitives, ensuring that performance scaling never compromises semantic fidelity.

As a consequence, the experiments documented in TVB-O are immediately available for gradient-based fitting, Bayesian calibration, or large sweeps over population parameters and stimuli, all while remaining auditable down to the original model specification. This closes the loop between ontology-backed description and optimization-ready execution, enabling side-by-side assessment of hypotheses and numerical backends under exactly matched scientific assumptions.

## 3. DISCUSSION

TVB-O is an open-source, collaborative knowledge base and software stack designed for large-scale brain simulations. Through an interactive GUI and programmatic API, it transforms high-level descriptions of computational modelling studies into multiple synchronized outputs: validated metadata records for humans and machines, simulator-ready code for multiple numerical backends, and provenance-aware reports. By standardizing how models, experiments, and results are specified, referenced, and shared, TVB-O provides a unified framework for reproducible digital brain twins. At its core, a human- and machine-readable representation captures local population dynamics of a nonlinear system, parameterization, network coupling, and observation models. RDF encodings supply persistent identifiers and cross-references that make curated resources easy to locate and cite, while templated exports bundle the exact software configuration so that derived artifacts remain directly accessible. Ontology-backed linkages expose the mathematical intent in a form that different simulators can interpret consistently, delivering interoperability across heterogeneous toolchains. Versioned specifications and provenance records ensure that published configurations can be reinstantiated and extended without ambiguity, promoting durable reuse of simulation data. By integrating a computational ontology, a knowledge graph, and code-generation backends, TVB-O accelerates hypothesis development and bridges disciplinary boundaries in computational neuroscience.

All mathematical equations are stored in a unified symbolic form independent of any programming language or simulator. This abstraction has profound implications: the same specification simultaneously drives simulation code generation across multiple backends (Python, Julia, JAX, C, LEMS), analytical derivations, human-readable LaTeX rendering for reports, and automated consistency checking. When a parameter is renamed, specific values are changed or an equation refined, changes propagate automatically to all downstream artifacts, namely the execution code, documentation, and analytical tools, remaining synchronized without manual re-implementation. This eliminates the endemic problem of divergence between “what the paper says,” “what the code does,” and “what the documentation claims,” ensuring that mathematical intent is preserved across translations. The symbolic layer also enables operations impossible with pre-compiled models: users can programmatically query dependencies (“which parameters affect excitatory gain?”), construct derivative quantities, verify dimensional consistency, and generate continuation programs for bifurcation analysis - all from the same source. This representation establishes equations as first-class, queryable entities in the knowledge graph rather than opaque procedural code.

What is new in TVB-O is a coherent contract that links semantic description, execution, and reuse. An OWL-based vocabulary makes core modeling assumptions explicit (for example, the structure of coupling, the handling of delays, and the mapping from model states to observations), reducing ambiguity across notations. Typed specifications allow models to be extended or recombined without bespoke glue, so dynamical systems, stimuli, networks, and observation pathways remain interchangeable within a consistent schema. A curated, versioned library binds models and studies to persistent identifiers that support reproducible reinstantiation and systematic comparison. An accessible browser and guided builder enforce consistency while lowering the barrier to configuration. A templated code-generation stack renders the same symbols and units to multiple numerical backends, keeping human-readable reports, machine records, and executables synchronized from a single source. Finally, a direct bridge to optimization workflows exposes ontology-defined parameters as search spaces for gradient-based fitting and large-scale sweeps while preserving provenance. Together, these elements replace ad-hoc conventions with a principled, interoperable workflow and provide a fair basis for reuse, cross-backend benchmarking, and parameter inference.

### 3.1. Limitations and Strengths of the TVB-O Framework

TVB-O addresses critical gaps in reproducibility and interoperability for whole-brain modeling, yet several limitations shape its current scope and utility.

#### 3.1.1. Curation scalability

The database currently contains 12 curated studies and 78 models - a meaningful but incomplete sample of published work. Manual annotation from literature remains labor-intensive, creating a bottleneck as the field expands. Each entry requires careful extraction of parameters, equations, and metadata from heterogeneous source formats (prose, tables, code), followed by validation against the schema. As modeling approaches diversify and publication rates increase, maintaining comprehensive coverage will strain expert curation capacity.

The LinkML schema and validation pipeline support distributed community contribution. Future integration of large language models for automated metadata extraction from PDFs could accelerate curation while maintaining quality through expert review of machine-generated candidates. The versioned repository structure allows incremental expansion without disrupting existing entries.

#### 3.1.2. Biological resolution

TVB-O deliberately targets mean-field population models tractable for whole-brain simulation, abstracting away ion channels, dendritic computation, and synaptic plasticity. This design prioritizes compatibility with neuroimaging observables and computational efficiency but limits applicability to subcellular hypotheses. Cellular heterogeneity within populations, detailed receptor kinetics, and morphological features are not represented at the resolution captured by standard neural mass formulations, potentially constraining mechanistic interpretation of certain disease processes or pharmacological interventions.

The framework’s modular structure accommodates future biological detail: biophysically detailed neuron models can be incorporated as alternative local dynamics classes, and cross-scale mappings to resources like NeuroML would enable hybrid multiscale workflows without redesigning core semantics. Gene Ontology and KiSAO alignments already provide scaffolding for richer biological annotation as experimental techniques advance.

#### 3.1.3. Backend dependencies

Code generation relies on external simulators (TVB, Julia, JAX, C) whose maintenance and API stability lie outside our control. Breaking changes in target frameworks require updating translation templates, and numerical precision differences across backends can introduce small divergences in long simulations. If a supported framework undergoes deprecation or fundamental redesign, corresponding export pathways may break until templates are revised, temporarily limiting user options.

Because the symbolic specification remains stable across backends, the template architecture enables adding new translators without altering curated models. Community-contributed backends and ongoing benchmarking will help maintain coverage as the simulator ecosystem evolves. JAX’s JIT compilation offers a pathway toward hardware acceleration for parameter-intensive workflows, and emerging GPU-native solvers can be integrated by adding targeted templates. For example, TVB-O supports the creation of CUDA code comparable to the one generated by RateML [61] and is therefore interoperable with external workflows.

#### 3.1.4. Validation coverage

While TVB-O enforces internal consistency (schema validation, unit checking, provenance linking), it does not automatically verify that generated code reproduces quantitative results from original publications. Ambiguities in source descriptions, like missing parameters, conflicting values across figures and tables, or partial reports, can necessitate conservative annotation of uncertain values. Users must independently confirm that reinstantiated experiments match published benchmarks, as systematic regression testing against original time series or bifurcation diagrams is not yet automated at scale.

Making validation gaps explicit rather than silently imputing defaults strengthens scientific integrity. Planned integration of automated regression testing against published benchmarks will systematically flag discrepancies for curator review, building confidence incrementally. Conservative annotation of uncertain values preserves the distinction between reported and assumed parameters, supporting meta-analyses of reporting practices. With the ontological integration, missing values are assumed from the original model publications.

#### 3.1.5. Adoption barriers

Effective use requires familiarity with dynamical systems, ontology structures, and YAML metadata, a learning curve for researchers accustomed to script-based workflows or point-and-click simulators. Understanding coupling semantics, observation models, and provenance graphs demands conceptual investment. Integrating TVB-O into established laboratory pipelines requires adapting existing code, retraining collaborators, and reconciling ontology-backed specifications with legacy data formats. These are transition costs that may delay uptake in resource-constrained settings or groups with already established computational pipelines.

The graphical user interface lowers barriers for model discovery and configuration without programming. Comprehensive documentation, worked examples in multiple domains, and integration with EBRAINS infrastructure reduce onboarding complexity. As community adoption grows, shared teaching materials and domain-specific tutorials will further accelerate uptake. The symbolic representation itself aids pedagogy: students can inspect dependency graphs and equation forms directly rather than reverse-engineering implementations.

#### 3.1.6. Real-time constraints

The framework optimizes reproducible research workflows rather than low-latency applications. Compilation overheads, particularly for the TVB pathway or un-JIT-compiled backends, may exceed real-time requirements for closed-loop clinical systems, intraoperative decision support, or brain-computer interfaces. Deterministic scheduling, hardware interfacing, and safety certification needed for time-critical medical applications lie beyond the current scope.

Persistent identifiers, provenance graphs, and machine-readable metadata align with journal and funder mandates for open science. The framework demonstrates immediate value for literature synthesis, cross-study comparison, and parameter exploration, practical benefits that drive adoption independent of real-time performance. Symbolic specifications also facilitate long-term preservation: equations remain interpretable even if specific simulator versions become obsolete.

In summary, TVB-O’s limitations reflect deliberate design choices (population-level scope, research-oriented workflows) and resource constraints (manual curation, backend coverage). Its symbolic, modular, open architecture and alignment with FAIR principles position the resource to evolve alongside computational neuroscience needs through community engagement and methodological advances in automated knowledge extraction.

### 3.2. Comparison with existing modeling tools and metadata initiatives

#### 3.2.1. Ontologies, knowledge bases, and metadata frameworks

TVB-O enhances initiatives like NeuroML and related specification languages by focusing on mean-field and whole-brain network models and combining formal semantics with executable artifacts. While NeuroML emphasizes cellular to network-level components, TVB-O provides a domain ontology, metadata schema, and translators aimed at large-scale network simulations and analysis workflows. These efforts are complementary rather than competitive, and interoperability remains a key goal.

Beyond format alignment, TVB-O links diverse parameter nomenclatures to shared ontology classes, enabling disambiguation without enforcing unification: different notations stay intact but refer to the same concept. For each annotated neural-mass model, TVB-O can produce LEMS XML descriptions; these artifacts can be used by RateML to generate Python or CUDA kernels, advancing interoperability toward GPU-based simulators without rewriting model logic.

#### 3.2.2. Computational neuroscience software

TVB-O interacts with simulators and analysis libraries by translating a single semantic specification into backend-specific executables and metadata. In practice, it serves as an extensible middle layer that aligns model intent with the numerical capabilities of individual frameworks: the same experiment description can be rendered as C, Julia, JAX, or Python. Currently, TVB-O supports the syntax of TVB, Julia’s NetworkDynamics.jl [62], and TVB-Optim [60], for efficient model computation and parameter optimization. The template system allows new engines to be added with minimal effort. The same templated contract also pushes capabilities back into those ecosystems. For example, registering a new neural mass model in TVB-O instantaneously generates the boilerplate required to load it inside TVB or other supported backends, keeping syntax and semantics synchronized without bespoke glue code. Benchmarks over this shared contract (see Supplement for the runtime benchmark figure) confirm that TVB-O maintains the characteristic performance hierarchy of native pathways: C fastest, Julia and un-jitted JAX closely aligned, Python slower, and TVB incurs higher orchestration costs, while ensuring that identical equations are compared fairly. Rather than competing with other simulation frameworks, TVB-O delegates execution to them, providing the semantic “mediator” that keeps results interoperable and reproducible across heterogeneous toolchains [26]. Additionally, from the same symbolic representation, TVB-O automatically creates continuation programs for automated bifurcation analysis, enabling model exploration and transfer of dynamical features across models, such as aligning excitability regimes among different model families.

### 3.3. Implications for adopting FAIR practices in computational neuroscience

TVB-O serves as a knowledge interface linking biology, mathematics, and computing by aligning controlled vocabularies and metadata with executable specifications, ensuring unambiguous communication and enhancing reproducibility [63]. Its GUI, model builder, and multi-language code generators lower entry barriers and foster cross-disciplinary collaboration by providing streamlined access to semantically rich, reusable models and experiments - much like ModelDB [64] - but with explicit semantics and platform independence. Integration with the EBRAINS Knowledge Graph [9] enables shared, centralized access to annotated models and data, while ontology-backed metadata enhance search and discovery without centralized oversight. TVB-O simplifies research workflows, from literature mining to model building, code generation, simulation, analysis, and sharing, with comprehensive documentation and examples available for all users regardless of discipline. Semantically typed specifications and translators enable seamless exchange and interoperability with tools such as NeuroML, TVB, and Julia-based libraries, promoting multiscale co-simulations between simulator platforms [4] and supporting Semantic Web principles [33] for integration with domain ontologies. Standardized, provenance-aware descriptions of models and experiments support replication and reuse across platforms; as the knowledge base grows, new evidence can be incorporated while preserving versioned specifications, strengthening the foundation of cumulative scientific research. Collectively, this provenance-traced workflow aligns computation with the same semantic contract and makes comparisons auditable, enabling fair computational research that treats models, analyses, and benchmarks on equal footing while increasing the share of studies that remain findable, accessible, interoperable, and reusable (FAIR).

#### 3.3.1. Optimization-ready workflows extend the reach of TVB-O

The tight coupling between TVB-O and TVB-Optim ensures that ontology-curated experiments can transition directly into gradient-based inference, Bayesian calibration, or high-throughput parameter sweeps without bespoke code. Because the optimizers operate on the same identifiers, units, and provenance captured during curation, parameter updates and observation metrics remain interpretable to neuroscientists who are chiefly interested in the biological hypotheses rather than implementation mechanics. This closes the gap between semantic description and numerical experimentation, giving researchers a reproducible path from curated knowledge to quantitative fitting.

By aligning differentiable execution with FAIR metadata, the integration also expands how results can be shared, audited, and reused. Laboratories can publish the ontological specification together with optimization trajectories, confident that other groups can replay the analysis on heterogeneous hardware or substitute alternative solvers without changing the scientific contract. In practice, this lowers the barrier to hypothesis testing across sites, accelerates benchmarking of emerging simulators, and strengthens the evidential weight of claims built on optimization-driven studies.

#### 3.3.2. Model selection and cross-model reasoning

Multiple neural mass models can produce similar macroscopic phenomena, such as resting-state-like activity, which raises a practical question about selecting the right model for a specific hypothesis. By presenting model semantics, parameter regimes, and literature-based usage side by side, TVB-O assists researchers in choosing an appropriate NMM for the phenomenon and translating parameterized hypotheses across models without altering their meaning.

#### 3.3.3. From empirical knowledge to executable hypotheses

Many variables in a brain network model serve as natural interfaces to empirical measurements and biological pathways. Instead of striving for exact replication of specific datasets, TVB-O promotes mechanistic alignment: include features relevant to the question and connect them to multimodal evidence (electrophysiology, imaging, genetics) where appropriate. Because parameters and states are typed and linked to definitions, the same biological perturbation can be consistently mapped across models. For example, implementing an excitation-inhibition shift by adjusting an inhibitory time constant in Jansen-Rit and identifying the corresponding inhibitory handle in another model through the ontology. In practice, this enables systematic hypothesis creation: starting from an observed effect or a biological pathway, the ontology surfaces candidate parameter handles and expected model-level consequences, making it straightforward to formulate and prioritize testable “what-if” perturbations. This approach makes it possible to incorporate semantic knowledge bases and measured effects into executable, hypothesis-driven simulations while maintaining comparability across different model families.

### 3.4. TVB-O as an educational resource for computational neuroscience research and teaching

Brain modeling is naturally interdisciplinary. TVB-O reduces barriers to entry by linking formal definitions with practical tools. The clear connection between ontology concepts, equations, and executable code encourages hands-on learning and creates a shared language for biology, physics, mathematics, and computation.

#### 3.4.1. Future Directions for ontology-driven brain modeling and simulation

We will expand literature coverage and strengthen semantic integration with domain ontologies (e.g., Gene Ontology [40], Human Phenotype Ontology [65], Cell Ontology [66], Uberon [49], NeuroML [23]) to support multiscale models [4] and closer links to empirical data. Graph-embedding methods will be explored for link prediction and hypothesis generation [67]. We also envision ontology-guided LLM workflows via retrieval-augmented generation (RAG) and knowledge grounding to improve transparency and reduce hallucinations in documentation and curation tasks [68]. Roadmap items include enhanced fitting pipelines, automated sensitivity and bifurcation analyses, and expanded backend support.

### 3.5. Conclusion and outlook

TVB-O formalizes large-scale brain modeling into a structured, interoperable, and executable framework. By standardizing model and experiment descriptions, it simplifies the configuration and execution of simulations and promotes systematic, FAIR research practices. As the resource evolves, we expect broader community adoption, improved reproducibility, and closer integration of models and data across scales. We encourage data repositories and journals to accept simulation metadata alongside results to enhance accessibility, searchability, and traceability. While focused on whole-brain simulations, the approach can be extended to annotate and execute dynamical-systems models beyond neuroscience.

## Data availability

All data supporting the findings of this study are available within the paper, its Supplementary Information, and the referenced public repositories.

Structural group connectomes used in this work include: (1) 32 Adult Diffusion MGH-USC HCP subjects with GQI fiber tracking (v1.2) [52, 53]; (2) 85 PPMI PD-patients with GQI fiber tracking (v1.1) [54, 44]; and (3) dTOR-985, a large normative connectome of 985 HCP subjects with GQI fiber tracking [42], available on Figshare (doi:10.6084/m9.figshare.c.6844890.v1). Connectomes (1) and (2) are available at https://www.lead-dbs.org/helpsupport/knowledge-base/atlasesresources/cortical-connectomes/.

Standard-space template atlases were retrieved from TemplateFlow [55]. The Human Connectome Project (HCP) Young Adult dataset (WU-Minn Consortium; PIs: David Van Essen and Kamil Ugurbil; 1U54MH091657), funded by NIH Blueprint institutes and the McDonnell Center for Systems Neuroscience at Washington University, provided the original data used for connectome construction; access is available at https://www.humanconnectome.org/study/hcp-young-adult/document/1200-subjects-data-release. Any additional curated derivatives used to generate figures are reproducibly produced by the code provided and described below.

## Code availability

All code, data model definitions, schemas, and the ontology are available at https://github.com/virtual-twin/tvbo. Documentation is available on the corresponding GitHub Pages site. Container build files (Docker) are included to support reproducible environments. A public web GUI is accessible at https://tvbo.charite.de/.

## Acknowledgements

This work used conversational AI assistants (Chat GPT-5 by OpenAI and Claude Sonnet 4.5 by Anthropic) to support manuscript editing and code scaffolding. All outputs were reviewed and validated by the authors, who take full responsibility for the content. P.R. acknowledges support by EU Horizon Europe program Horizon EBRAINS2.0 (101147319), VirtualBrainTwin(101137289), EBRAINS-PREP101079717, AISN101057655, EBRAIN-Health 101058516, EIC grant PHRASE 101058240, by the Digital Europe Programme TEF-Health (101100700), Shaiped (101195135), CoordinaTEF (101168074) German Research Foundation SFB 1436 (project ID 425899996); SFB 1315 (project ID 327654276); SFB 936 (project ID 178316478); SPP Computational Connectomics RI 2073/6-1, RI 2073/10-2, RI 2073/9-1; DFG Clinical Research Group BECAUSE-Y 504745852, Berlin University Alliance OpenMake, the Virtual Research Environment at the Charité Berlin and EBRAINS Health Data Cloud and the Berlin Institute of Health and Foundation Charité. P.R. and J.M. acknowledge additionally support by the Deutsche Forschungsgemeinschaft (DFG, German Research Foundation) - Project-ID 424778381 - TRR 295.

## 4. METHODS

### 4.1. Building a minimal, unambiguous vocabulary and logical axioms for brain simulations

#### 4.1.1. Mathematical framework

The ontology formalizes the mathematical structure underlying whole-brain network simulations. At the core, brain dynamics are described by coupled differential equations where the state *S_i_* of region *i* evolves according to:

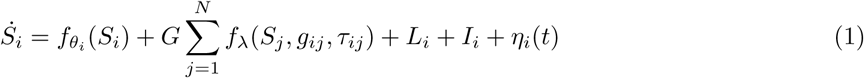

This expression captures five essential components: (1) **local dynamics** *f_θ_* (*S_i_*) governed by neural population parameters *θ_i_*; (2) **global coupling** that aggregates delayed inputs from distant regions *j* via white-matter connections with weights *g_ij_* and delays *τ_ij_*, scaled by gain *G*; (3) **local (surface) coupling** *L_i_* from geometrically neighboring vertices; (4) **external inputs** *I_i_* representing targeted stimuli; and (5) **stochastic terms** *η_i_* for noise. Delays derive from tract lengths *D_ij_* and conduction velocity *v* as *τ_ij_* = *D_ij_/v*.

The coupling function *f_λ_* transforms and sums afferent activity with explicit pre-synaptic, summation, and post-synaptic steps, preserving ordering semantics. Local coupling for surface-based simulations involves a spatial kernel *K*(*d_kl_*) operating over cortical distances. Full derivations and component-wise equations are provided in the Supplementary Introduction (Building Blocks of a Brain Network Model).

#### 4.1.2. Ontology structure and coverage

TVB-O is an OWL domain ontology (base IRI: http://www.thevirtualbrain.org/TVB-O/) that formalizes the core entities required to specify and execute whole-brain network simulations. Classes form a directed acyclic hierarchy, with optional multiple inheritance, defined principally by rdfs:subClassOf relations; OWL restrictions provide class-level constraints where needed. Authoring and programmatic access are supported through WebProtégé and Owlready2.

##### Scope

The ontology covers the five elements in Equation 1: (1) brain networks represented by structural weights *W_ij_* (equivalent to *g_ij_*) and tract lengths *D_ij_*; (2) local neural dynamics with state variables *S_i_* and parameters *θ_i_*; (3) coupling functions *f_λ_* operating on delayed states with *τ_ij_* = *D_ij_/v*, scaled by global gain *G*; (4) numerical integration; and (5) external perturbations *I_i_* and optional noise *η_i_*.

##### Mathematical representation

Equations are first-class entities: both left- and right-hand sides (and optional conditionals) are stored in LaTeX and in a plain symbolic form referencing only {*S_i_, θ_i_, W_ij_, τ_ij_, G, I_i_, η_i_*}. The Coupling class decomposes transformations into pre-synaptic processing, weighted summation over afferents, and post-synaptic transforms, preserving the order by which delayed inputs contribute to d*S_i_/*d*t*.

##### Semantics and validation

Axioms link symbols to their roles (has_state_variable, has_parameter, has_derivative), enabling automated consistency checks and cross-model comparison. All entities carry persistent URIs with SKOS annotations (labels, definitions, notes), and provenance follows W3C PROV. Coverage spans region- and surface-based models; connectomes provide *W_ij_* and *D_ij_*; observation models and exemplar neural mass families (e.g., Jansen-Rit, Epileptor, Larter-Breakspear, Hopfield) retain equations aligned with simulator conventions.

#### 4.1.3. Ontology population from TVB source code

From The Virtual Brain source code, we parsed each model to extract state variables, parameters, and functions; normalized and structurally analyzed function bodies; detected time derivatives; and mapped algebraic dependencies. Parsed elements are instantiated as ontology entities with axioms capturing math-ematical structure (has_parameter, has_state_variable, has_derivative, is_coefficient_of) together with model-specific annotations (equation text, definition, primary reference). To support reasoning and reuse, implicit runtime notions are made explicit (TimeDerivative, Function, DerivedVariable, ConditionalDerivedVariable), and required superclasses are introduced (e.g., GlobalConnectivity as a subclass of Network). Class-level constraints such as “a WeightsMatrix about some Network” (obo:IAO_0000136) are recorded where appropriate.

#### 4.1.4. Biological grounding via Gene Ontology

To link biological knowledge to model components, we curated vertebrate-relevant Gene Ontology processes [40] and mapped them to TVB-O entities using transparent rules (e.g., excitation/inhibition, excitation-inhibition balance, coupling, kinetics). Where supported by the literature, we asserted parameter-level links; for example, “negative regulation of AMPA receptor activity” (GO:0061907) is mapped to the r_NMDA parameter in the Larter-Breakspear model.

#### 4.1.5. Interoperability with Other Ontologies

We imported the Brain Atlas Ontology Model (AtOM) [39] from the SciCrunch NIF-Ontology repository (atlas branch). Additional details are provided in the Supplement (Integration of the AtOM ontology for brain atlas metadata).

### 4.2. Creating a compact ontology-aligned metadata schema for reproducible simulations

We designed a concise, ontology-aligned metadata model to describe brain network models and simulation studies in a simulator-agnostic manner. The model is specified in LinkML, which provides a readable schema and machine-enforceable types, and it is grounded in persistent identifiers drawn from the ontology for brain simulation (TVB-O) and from the AtOM atlas model. The guiding principle is to capture precisely those elements that determine a simulation outcome while keeping the structure compact enough to be authored and understood by researchers.

The schema organizes information around a small number of scientific concepts. Local population dynamics are described as named state variables governed by symbolic equations, accompanied by tunable parameters and optional derived quantities (for example, outputs or intermediate expressions). Each quantity may carry a numerical domain (lower and upper bounds and, where relevant, a sampling step) and a unit, allowing both validation and meaningful comparison across studies. Network structure is represented by a connectome that comprises a weights matrix and a tract-lengths matrix, linked to a parcellation whose regions are defined within a specific brain atlas and coordinate space. Interactions between regions are controlled by an explicit coupling term; numerical integration may include stochastic components (noise) with clearly declared parameters. External perturbations are modeled as stimuli targeted to specific regions and variables. Observation choices (for example, raw activity, BOLD, EEG/MEG) are recorded as monitors, making explicit which signals are produced by an experiment.

Two container levels provide context and provenance. A simulation experiment assembles a particular configuration-local dynamics, network, coupling, integrator, stimulus, and observation choices-into a complete run specification. A simulation study groups one or more experiments under a shared citation and metadata, enabling within-study comparisons and sensitivity analyses. Software environment and dependency information can be included to ensure that identical configurations can be recreated beyond the original computing setup; dataset and subject descriptors link experiments to empirical resources when applicable.

Equations are preserved in a symbolic form that is independent of any programming language. Left- and right-hand sides can be recorded, as can conditional expressions, together with optional LaTeX renderings. This representation supports multiple downstream uses: exact reproduction of the model specification, side-by-side comparison of formulations across studies, and automated generation of simulator-specific code or human-readable reports without drifting from the source description.

In practice, the LinkML schema yields both a human-readable template for authors and a rigorous, machine-checkable contract for tools. Metadata authored from the literature or through the GUI are validated against the schema before being integrated into the knowledge base, exported as FAIR records, or transformed into executable configurations for different simulation backends. By aligning fields with ontology identifiers from TVB-O (simulation concepts) and AtOM (atlas concepts), entries interoperate with the Semantic Web and remain unambiguous across datasets and software.

### 4.3. Creating a curated, versioned model database for retrieval, comparison, and reuse

#### 4.3.1. What is extracted, and how is the process done?

We curate a compact database of simulation studies and canonical models by translating the essential elements of each publication into an ontology-grounded, LinkML-validated description. Each entry names the neural population model and preserves its symbolic equations, states, parameters, and derived variables together with units and typical numerical domains. Network specifications are recorded as a parcellation linked to an atlas entity, a weights matrix, and tract lengths; coupling semantics, numerical integration settings (including stochastic terms), external stimuli, and observation choices are captured explicitly. Studies are organized as containers with one or more experiments so that within-study parameter sweeps and sensitivity analyses can be represented without duplicating common context such as the citation or software environment.

Annotation proceeds from the literature or companion code to a schema-checked YAML record. During curation, symbols in equations and parameter tables are resolved to stable ontology identifiers (from TVB-O and AtOM), synonyms are normalized, and units are harmonized; LinkML validation enforces types and required fields and rejects inconsistent or ill-typed assignments. Where publications supply algebraic expressions, we store left- and right-hand sides in a normalized symbolic form, enabling downstream rendering to LaTeX, code generation, and analytical checks from the same source. Provenance is retained through a citation key that maps to the shared bibliography and through software requirements that pin critical dependencies for reproduction.

The resulting records are versioned in the repository. Study entries reside under tvbo/data/db as YAML files, with a shared bibliography at tvbo/data/db/tvbo-literature-db.bib. Canonical model descriptions used across studies are kept alongside under tvbo/data/db/Model. To make the editorial state transparent, entries under active review are placed in tvbo/data/db/to_adjust!, and rejected or didactic examples are kept in tvbo/data/db/invalid. Accepted entries are compiled into a consolidated index (tvbo/data/db/db.pkl) for fast programmatic access. This layout supports incremental growth of the database while keeping the distinction between curated, provisional, and deprecated content clear.

#### 4.3.2. Handling missing parameter values not reported in a study

We prioritize faithfulness to the source over forced completeness. When articles omit values that affect reproducibility, we record the field explicitly as missing rather than inventing a number, and annotate the reason where known (for example, “not reported,” “fixed but unspecified,” or “visually estimated”). If a plausible default is required to run an example, we borrow values from the canonical formulation of the same model and state this as an assumption in the entry, leaving the original field empty so the distinction between reported and assumed values remains visible. Units and parameter domains follow the canonical model unless the study reports otherwise; any transformations applied during harmonization (for example, ms to s) are documented in the metadata.

Ambiguities are resolved conservatively. Conflicting values across figures, tables, or code are both recorded with a short note and the entry is flagged for review in to_adjust!/ until the discrepancy is settled. Where only a derived quantity is provided (for example, a composite coupling gain), we encode the equation and tag the underlying components as unknown rather than back-solving without explicit justification. This policy preserves the integrity of the database, enables transparent comparison across studies, and still allows complete, executable specifications once conservative defaults are supplied for illustration or benchmarking.

### 4.4. TVB-O Software Package

#### 4.4.1. Software dependencies and containerization

TVB-O is distributed as a Python package with an accompanying container that pins all dependencies for reproducible execution across platforms. The container bundles the ontology client, metadata validation, code generation, and simulation backends; optional accelerators (such as just-in-time compiled kernels) can be enabled but are not required. A single build step produces a ready-to-run image for the figures and benchmarks reported here.

#### 4.4.2. How the ontology is represented and accessed

The software exposes the ontology as a searchable knowledge base: concepts, relations, and annotations can be browsed in the GUI or queried programmatically. OWL entities (for example, NeuralMassModel, StateVariable, Coupling) are mapped to human-readable labels with stable identifiers. Users can search by exact names, synonyms, or symbols and expand parents and children to explore context. For interactive analysis, the same content can be viewed as a graph, exported to JSON, or summarized in tables. This unified interface supports both exploratory work and automated pipelines, such as creating a study specification from a selected set of ontology nodes.

4.4.3. *Symbolic model representation for platform-independent configuration*

Mathematical expressions for local dynamics, coupling, stimuli, and observation models are represented symbolically to support platform-independent configuration, LaTeX rendering, code generation, and dynamical-systems analysis (for example, Jacobians and fixed points). Symbolic forms serve as the single source for translators targeting different computational backends.

When a model is loaded from the knowledge base or a metadata file, its equations are normalized into a consistent symbolic form, with explicit left- and right-hand sides and optional conditions. Symbols are resolved against ontology labels and synonyms to avoid ambiguity; parameter units and typical ranges are retained where available. Dependencies between intermediate expressions are ordered automatically, allowing the same specification to drive code generation, analytical derivations, and human-readable reports without divergence.

#### 4.4.4. Templating for different backends

From a single specification, TVB-O emits simulator-ready code and standard descriptions. Templates generate classes for The Virtual Brain, lightweight Python functions, and Julia scripts; LEMS export produces XML that validates with the reference toolchain. A small set of normalization rules-for example, systematic handling of conditional expressions-ensures that equations execute consistently across targets while preserving their original mathematical meaning.

#### 4.4.5. Framework for executing simulations

Simulation runs are assembled from five building blocks: local dynamics; network connectivity, including delays derived from tract lengths; inter-regional coupling; numerical integration, with optional stochastic terms; and observation choices. TVB-O computes the required history window for delayed interactions, configures monitors (for example, raw activity or BOLD-like signals), and then executes the model either within The Virtual Brain or with a lightweight runtime suitable for rapid prototyping. The same experiment can be exported as code, ensuring that what is inspected in the interface is exactly what is executed.

Internally, simulations are represented as directed acyclic graphs of operations, including integration steps, coupling, stimuli, and observation models. This design aligns naturally with provenance capture, supports time-varying inputs, and enables deterministic replay and post-hoc derivations from intermediate results.

### 4.4.6. Framework for automated bifurcation analyses

From the same symbolic equations used for simulation, TVB-O generates scripts for continuation and bifurcation analysis. The system supports Julia’s BifurcationToolkit and a classic AUTO/NumCont workflow; both are produced directly from the model specification so that parameter names and equations remain consistent with the simulation. For reproducibility, a containerized runner is provided for the AUTO/NumCont path.

#### 4.4.7. Model reports

Parameter tables and equation summaries are rendered automatically from the same specification used to generate code. Outputs include human-readable Markdown/LaTeX and optional PDFs, ensuring that documentation and execution stay synchronized.

#### 4.4.8. Graphical user interface: architecture and implementation

The graphical user interface exposes the knowledge base, allows users to assemble valid combinations of model components, and exports the resulting specification as runnable code and FAIR metadata. Behind the scenes, a small set of services handle parsing of literature and metadata, ontology-backed search and graph navigation, code generation (including LEMS), execution, and visualization. The same services are available via a programmatic API for scripted workflows.

#### 4.4.9. Model configuration workflow

TVB-O turns conceptual model choices into an executable, FAIR specification through a short, guided workflow. Each step is validated against the schema and anchored to persistent identifiers so terminology remains unambiguous across tools and studies.

Researchers begin by selecting local population dynamics from the curated library or by importing a custom description. State variables, parameters, and equations can be inspected and adjusted as needed; equations remain in symbolic form so the same description supports analysis, code export, and reporting from a single source.

The structural network is then defined by choosing a parcellation and a connectome comprising weights and tract lengths. TVB-O derives biologically plausible delays automatically from tract lengths and a conduction velocity, while subject- or cohort-specific matrices can be supplied to match empirical data. Region labels remain aligned to atlas entities for comparability across studies.

Inter-regional coupling and external inputs are specified next. The coupling function and its parameters are chosen with explicit pre- and post-synaptic semantics that clarify how signals are transformed and aggregated across the network. Targeted stimuli can be attached to specific regions and state variables; inputs appear as first-class terms in the equations.

Numerical integration and observables are configured by selecting a step size and deterministic or stochastic schemes, and by choosing observation outputs such as raw activity or BOLD-like signals. Sampling periods, units, and monitor settings are harmonized with the model to support meaningful comparisons and downstream analysis.

Finally, the experiment can be executed directly or exported as simulator-ready code, LEMS XML, FAIR metadata, and a human-readable report. The single specification captures provenance and software requirements, enabling faithful reproduction across systems and backends without re-implementation.

### 4.5. Optimization workflows - TVB-Optim interface

The optimization workflow begins by requesting a JAX export of a curated simulation experiment. TVB-O assembles a complete simulation state that captures ontology-linked initial conditions, the connectome, coupling, integrator settings, and observation pathways, and it emits a compiled kernel whose numerical implementation matches the curated equations. During export, the helper configures floating-point precision (single or double), harmonizes monitor choices (for example, optional FFT-based BOLD convolution), and serializes all arrays so they are ready for accelerator execution without manual rewrites.

TVB-Optim wraps this state in lightweight parameter classes that behave like JAX arrays while retaining bounds and scaling. By registering these classes as pytrees, the framework partitions the state into differentiable (free) and static components, detaching quantities such as structural connectivity while exposing the parameters slated for fitting. Normalization and clipping occur transparently, so constraints specified in the metadata are enforced throughout optimization.

Systematic exploration is achieved by assigning axes to any ontology-referenced parameter. Grid, uniform, or NumPyro-derived axes generate value sets that are automatically broadcast to the correct tensor shapes, and the resulting collections are reshaped for vectorized execution using JAX’s vectorized mapping or for distributed evaluation using its parallel mapping. TVB-Optim provides a deterministic sequential runner for iterating through parameter combinations and a parallel engine that batches them across devices while recombining static and variable components on the fly.

Loss functions operate on the same simulation state structure, drawing on observation utilities (for example, functional connectivity correlations or root-mean-square error) that accept time-series outputs. Gradients are obtained via automatic differentiation and supplied to Optax optimizers; callback hooks provide logging, early stopping, and best-of-run checkpoints. Throughout, the state retains the identifiers and provenance supplied by TVB-O, ensuring that optimization outputs can be returned to the knowledge base without loss of context.

4.5.1. *Stimuli and observation models*

Stimuli can be defined analytically (for example, pulses or sinusoids) or loaded from data (for example, an audio file resampled to the simulation step). They are targeted to regions and specific state variables and are treated as first-class inputs in the equations. Observation models map hidden states to observable signals through configurable transfer functions and can be combined into simple pipelines, such as state selection followed by a hemodynamic transform. Both are described symbolically and therefore export alongside the rest of the model.

### 4.6. METHOD for use cases: Study configurations, stimuli, and observation pipelines

Below, we outline the methods used in the use cases showcasing TVB-O’s semantic model specification framework to (i) compose an ontology-backed graph to map disease entities to model parameters; (ii) generate dependency graphs and parameter continuations with matched stimuli across models; (iii) declare observation pipelines as symbolic DAGs from simulated states to BOLD and derived features; and (iv) specify targeted, time-varying stimuli linked to atlas regions and state variables.

#### 4.6.1. Methods for Use case 1: Linking semantic knowledge sources to derive mathematical parameters from biological processes

We integrate a disease-focused knowledge graph with TVB-O by composing subgraphs from NeuroMMSig’s Biological Expression Language (BEL) resources with ontology-backed relations between TVB-O and the Gene Ontology into a single, analyzable network that links biological processes to model parameters. For reproducibility, BEL parsing is cached and fixed random seeds are used wherever layout or sampling routines are involved.

NeuroMMSig graph construction. We iterated over all BEL scripts distributed with the NeuroMMSig knowledge package, suppressed diagnostic messages from the BEL parser, and used a fixed seed (1312). Each BEL file was parsed to a PyBEL graph and stored in a cached list under the package’s temporary directory to avoid repeated input/output. We then constructed a multi-directed graph in NetworkX by traversing edges across all BEL subgraphs and mapping each BEL node to a canonical label using a dedicated label-mapping utility. Gene Ontology nodes were normalized by stripping the “GO:” prefix where present. Nodes were added with their inferred type and annotated with a source attribute (“NeuroMMSig”); edges between identical labels were ignored to prevent self-loops.

TVB-O to GO graph construction. We obtained the set of GO terms aligned to TVB-O (the descendants of a dedicated TVB-GO class) and built a TVB-O/GO graph from those terms. Next, for every ontology class and object property defined in TVB-O, we retrieved property annotations and added edges between the corresponding class labels, typing the edge by the property’s label. Nodes were labeled by origin-“GO” for Gene Ontology terms and “TVB-O” otherwise. The NeuroMMSig and TVB-O/GO graphs were then composed into a single multi-directed graph that preserves node and edge attributes from both sources.

Shortest-path linking from disease to model parameters. To illustrate how disease biology connects to model handles, we computed shortest paths on the undirected view of the composed graph between Alzheimer-related nodes (MeSH: Alzheimer Disease; HGNC: APP; CHEBI: amyloid-beta) and TVB-O model elements. For one panel we targeted the set of Jansen–Rit parameter classes retrieved programmatically from the ontology; for others we targeted exemplar classes (such as ReducedSetHindmarshRose) and individual parameters (such as b_JR). Paths from each source–target pair were merged into a subgraph for plotting. Before rendering, nodes corresponding to ontology classes were renamed to use their symbol where available (falling back to the label), and nodes were colored by origin-GO (teal), TVB-O (blue), or NeuroMMSig and other sources (orange). Layout used Graphviz’s dot with the root fixed at “Alzheimer Disease,” and edges were drawn with curved arrowheads for legibility.

Exports and filtering for visualization. Beyond the figure panels, we exported JSON serializations of the merged graph for interactive inspection. We first extracted the largest connected component, labeled each node with an origin tag (GO, NeuroMMSig, TVB-O) and a Boolean cross-link flag if it participates in at least one edge that bridges different origins, then sanitized all node and edge attributes for JSON compatibility. Two exports were provided: a whole-graph view and a filtered view that retains GO terms together with their first-order TVB-O/NeuroMMSig neighbors and those neighbors’ immediate connections. These artifacts enable downstream web visualization and reproducible sharing of the integration results.

#### 4.6.2. Methods for Use case 2: Comparing Neural Mass Models

We compared the Jansen-Rit and the reduced Wong-Wang (excitatory/inhibitory) models using a unified, ontology-backed workflow that generates dependency graphs, one-parameter bifurcation diagrams, targeted time series under identical stimuli, and two-dimensional parameter explorations. Models were loaded by their curated identifiers and executed via the Python backend to ensure deterministic results and consistent semantics across steps (see the accompanying script in the manuscript code directory).

For each model, we rendered the symbolic dependency graph of states, parameters, and derived variables and colored nodes by excitatory or inhibitory role using the curated annotation. Because the visualization is produced directly from the model specification rather than drawn by hand, edges reflect the algebraic dependencies used during simulation (Figure 5 **a,b**).

We then computed one-parameter bifurcation diagrams from the same symbolic equations to map qualitative regime changes. In the Jansen-Rit model, the control parameter *µ* (continuation input control) was varied from −0.1 to 0.4 and the variable of interest was *y*_2_; periodic orbits were enabled to detect limit cycles. In the reduced Wong-Wang model, the external input *I*_ext_ was varied from −0.1 to 0.1 and the variable of interest was *S_e_*; continuation step controls were set to d*s* = 10^−8^, d*s*_min_ = 10^−10^, and d*s*_max_ = 10^−2^ with verbose reporting. Legends summarize fixed points and limit cycles (Figure 5 **c,e**).

To ground the bifurcation view in trajectories, both models were stimulated with the same pulse train targeted to their designated stimulation variables (Jansen-Rit: *y*_4_; reduced Wong-Wang: *S_e_*). The stimulus had onset at 4000 ms, period *T* = 8000 ms, pulse width *τ* = 1000 ms, and amplitude 0.05. Each model was simulated for 8000 ms from zero initial conditions, the first 2000 ms were discarded as transient, and three representative traces spanning the bifurcation regimes were plotted. For Jansen-Rit we used *µ* ∈ {−0.1, 0.09, 0.2} and displayed state *y*_1_, marking the corresponding *µ* values on the bifurcation panel; for reduced Wong-Wang we used *I*_ext_ ∈ {−0.1, −0.05, 0.1} and displayed *S_e_*, setting *J_N_*= 0.5 for these examples (Figure 5 g-l).

Next, we performed two-dimensional parameter explorations on a 20 x 20 grid and computed summary statistics of the resulting trajectories to delineate operating regimes. For Jansen-Rit we swept *µ* over [-0.1, 0.4] and *C* over [1, 200] and plotted the mean of (*y*_1_ − *y*_2_) as a filled contour (Figure 5 **d**); for reduced Wong-Wang we swept *I*_ext_ over [-0.1, 0.1] and *J_N_* over [0, 1] and plotted the mean of *S_e_* analogously (Figure 5 **d**).

All simulations used the Python engine with a duration of 8000 ms per run; unless stated otherwise, other solver settings followed the model defaults. To make the grid search reproducible and efficient, summary statistics (mean, maximum, variance) were memoized to a local cache keyed by model name, parameter names and values (rounded to three decimals), and the queried state variable.

#### 4.6.3. Methods for Use case 3: Post-processing of simulated data with executable directed acyclic graphs

This use case implements observation models as directed acyclic graphs that map simulated neural states to BOLD-like signals and downstream features. An observation model wires function nodes-each with a symbolic expression, typed inputs, default parameters, and units-and executes them in topological order.

Input time series from the simulator are treated as labeled tensors with axes corresponding to time, state, region, and mode. Selectors restrict operations to specific state variables (for example, *V* or *W*) or region subsets. A shape-safety utility expands arrays as needed to four dimensions to avoid broadcasting ambiguities. Time can be passed explicitly when kernels depend on it.

The BOLD forward path comprises three steps. First, a hemodynamic response function is constructed as a Volterra or canonical kernel parameterized by amplitude and time constants and normalized to unit area. Second, state-wise convolution uses an FFT-based routine with zero-padding and trimming to preserve signal length; parameters such as mode, padding, and data type are fixed to ensure deterministic behavior across platforms. Third, optional down-sampling can align the output to a target repetition time while retaining the original simulation time base. Both raw and BOLD-transformed signals are returned as labeled time series.

Feature derivation is expressed as terminal nodes in the same directed acyclic graph. For functional connectivity derived from BOLD signals, regional time series are z-scored along time and pairwise Pearson correlations are computed (upper triangle reported). For comparison, an identical derivative on the raw signals is included. Derivatives consume upstream node outputs and are computed deterministically.

Implementation details. Functions and graphs are declared symbolically and rendered to numerical routines at execution time using numerical Python backends. Node metadata records package versions and parameters. The graph, with its inputs and outputs, is serialized with the simulation experiment, allowing alternative hemodynamic response functions or kernels to be substituted without altering the experiment description.

#### 4.6.4. Methods for Use case 4: Computational representation of complex stimuli paradigms

We instantiated a whole-brain network model using the 17-region Yeo2011 resting-state functional network parcellation [59] and the dTOR-985 normative structural connectome from [42], which aggregates diffusion MRI tractography from 985 Human Connectome Project subjects. The connectome provides region-to-region weights (streamline counts) and tract lengths; for display, weights were min-max normalized via (*W* − *W*_min_)*/*(*W*_max_ − *W*_min_) to emphasize relative connectivity strength, and the structural graph was rendered with atlas-based node colors. Tract lengths are reported in millimeters and used to compute conduction delays assuming a fixed velocity.

Local population dynamics were modeled with the Generic 2-Dimensional Oscillator (G2D), a phenomenological two-state (*V*, *W*) neural mass model that generalizes the FitzHugh-Nagumo system to reproduce a wider range of physiological dynamical regimes. The G2D evolves according to:

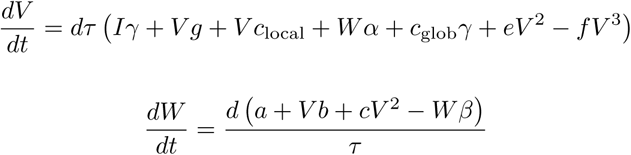

where *V* represents the fast excitatory variable and *W* the slow recovery variable. Parameters were set as follows: *a* = 0.001 (modified vertical shift of the nullcline to adjust baseline dynamics), *b* = −10.0 (linear slope of the configurable nullcline), *c* = 0.0 (parabolic term), *d* = 0.02 (temporal scale factor), *e* = 3.0 (coefficient of the quadratic term), *f* = 1.0 (coefficient of the cubic term), *g* = 0.0 (linear term coefficient), *α* = 1.0 (feedback scaling from *W* to *V*), *β* = 1.0 (self-feedback of *W*), *γ* = 1.0 (input current scaling), *τ* = 1.0 (time-scale hierarchy parameter), and *I* = 0.0 (baseline input). All remaining G2D parameters retained their canonical values as defined in the ontology.

Inter-regional coupling was configured via a linear coupling function with gain *a* = 0.5, which scales the weighted sum of delayed afferent activity from connected regions. Coupling acts on the fast variable *V* and propagates through the structural network with delays derived from tract lengths and conduction velocity.

An audio waveform was supplied as a time-varying external stimulus through the perturbation interface. The audio signal was internally resampled to match the simulation time step and truncated to the run duration (4000 ms). The stimulus was targeted to node 3 (ventral somatomotor network, encompassing primary auditory cortex) and injected into state variable *W* to modulate the slow recovery dynamics. Stimulus amplitude was scaled by a factor of 0.003 for visualization alongside neural time series.

We executed two simulation runs with identical initial conditions and parameters: one with the acoustic stimulus applied to node 3, and one control run without stimulation. Both simulations used a time step Δ*t* = 1 ms, total duration of 4000 ms, and integration via the default deterministic scheme. Time series from state variable *V* were extracted from all 17 regions and used to compute functional connectivity as the Pearson correlation coefficient matrix across regional time series. For visualization, the stimulated condition occupies the upper triangle of the functional connectivity matrix, while the lower triangle displays the resting (no-stimulus) condition for direct comparison.

