## Supplement for "The Virtual Brain Ontology: A Digital Knowledge Framework for Reproducible Brain Network Modeling"

---

*Keywords:* Brain Simulation, Ontology, Knowledge Base, Knowledge Engineering, Semantic Code Generation Framework, Scientific Computing, FAIR research

---

[†] The authors contributed equally to this work as co-last authors.

[*]Corresponding authors

### 1. Supplement

#### *1.1. SUPPLEMENTARY INTRODUCTION*

##### *1.1.1. Supplementary Table S1: Challenges, needs, and TVB-O solutions*

Table 1: Challenges, needs, and solutions provided by TVB-O for enabling FAIR large-scale brain modeling workflows.

| Category | Challenge | Need | TVB-O Solution |
|---|---|---|---|
| Model Specification | No standardized high-level language for networks of coupled nonlinear dynamical models. | Intuitive, editable, simulator-agnostic model definitions. | Declarative metadata schema for specifying brain dynamics and simulation schemes. |

| Category | Challenge | Need | TVB-O Solution |
| --- | --- | --- | --- |
| Semantic Consistency & Unification | Heterogeneous terminology, implicit assumptions, and absence of a domain-wide ontology for large-scale brain models. | Unified controlled vocabulary, semantic alignment, and metadata standard. | Computational ontology defining core entities and relations, linking mathematical models to biological interpretations with a controlled vocabulary and metadata schema. |
| Reproducibility | Incomplete experiment metadata or unclear formalism. | Reproducible annotation of modelling studies. | Executable, MIASE-compliant model specifications with provenance-aware metadata. |
| Model Reusability | Fragmented model landscape; difficult to compare, adapt or extend existing models. | Cross-compatible, adaptable model components | Curated repository with tools for integration and reuse |
| Code Portability | System-dependent code limits reuse | Platform independent workflows | Automated translation into simulator-specific code and other specification languages |
| Interoperability | Poor transfer of models across modelling platforms. | Unified interface for model translation. | Ontology-driven mapping of model components across simulators and integration with other ontologies. |
| Integration | Difficulty linking across biological scales | Conceptual framework allows multi-scale expansion | Ontology interoperable with biological knowledge graphs covering genes, neurons and networks. |
| Automation | Custom non-standardized and error-prone modelling pipelines | Scalable, efficient simulation workflows based on templating. | GUI and semantic engine for model querying and code generation |

Whole-brain network models (BNMs) approximate population dynamics across anatomically connected regions to explain measured signals and mechanisms of disease. When individualized with subject-specific data, these models act as “digital twins,” providing a mechanistic scaffold for simulating, analyzing, and ultimately personalizing interventions at the systems level. The Virtual Brain has catalyzed this field by offering a rich library of local population models and a neuroinformatics stack for building and simulating large-scale networks.

At the same time, the rapid growth of models, datasets, and software has created a fragmentation problem: results are difficult to find, integrate, and reuse because terminology, mathematical notations, and metadata practices vary widely. Reproducible research at scale requires contextualized, machine-actionable metadata that make assumptions, units, and dependencies explicit while preserving human readability. In practice, this means treating model components, workflows, and outputs as first-class, citable entities with stable identifiers and minimal yet sufficient metadata.

Knowledge graphs and computational ontologies address this need by providing a shared, formal vocabulary for domain concepts and their relations. Building on Semantic Web foundations (RDF/OWL) and standards for terminology and provenance (SKOS, PROV), ontology-backed resources enable software and readers to interpret “what a thing means,” not just how it is formatted. Existing efforts in neuroinformatics have shown the value of such standardization for atlases, neuroimaging results, and model descriptions; however, a unified namespace tailored to neural-mass and whole-brain network modeling has been missing.

TVB-O fills this gap. It formalizes the primitives of large-scale brain simulation (models, parameters, equations, networks, coupling, integrators, stimuli, observation models) and aligns them with a minimal metadata scheme and a provenance model, so that complete simulation studies can be specified, validated, queried, and reproduced. The same specification drives executable code generation and human-readable reports, ensuring that meaning, documentation, and execution remain synchronized. This supplement details how these ingredients come together in practice-demonstrating provenance-aware replication, curated normative connectomes, flexible observation models, and cross-backend benchmarking-while keeping the focus on clarity, comparability, and FAIR reuse.

###### *1.1.2. Building Blocks of a Brain Network Model*

The foundational principle of dynamic brain models is rooted in dynamical systems theory, which uses differential equations to represent the temporal evolution of a system's state based on the physical laws that govern it [1]. The change in brain activity or state $S$ of a region $i$ can be described by a general equation that accounts for local population parameters $\theta_i$ , the short-range input based on surface connectivity $L_i$, the coupled input from other distant network areas $C_i$ depending on their anatomical connectivity, and a global coupling scaling factor $G$ (equation adapted from [2]):

$$\dot{S}_i = \frac{dS_i}{dt} = f(S, \theta_i, L_i, C_i, G) \tag{1}$$

1. **Local neural population dynamics**: In an uncoupled and non-perturbed system, $\dot{S}$ describes how the activity of a neural population in a particular region changes over time, based on its previous state and a set of local population parameters $\theta_i$. The function $f_{\theta_i}$ incorporates the set of differential equations for a given neural mass model.

$$\dot{S} = f_{\theta_i}(S(t-1)) \tag{2}$$

2. **Global coupling**: Represents the long-range input from distant nodes via white-matter fibers, scaled by a global coupling factor.

$$C_i = G \sum_{j}^{N} f_\lambda(S_i, S_j, v, g_{ij}) \tag{3}$$

The term $C_i$ denotes the summed input from all other distant network areas $N$ to region $i$. The generic coupling function $f_\lambda$ with the parameter set $\lambda$ describes how the activity in region $j$ influences region $i$, taking into account the the anatomical connection strength $g_{i,j}$ between these regions. The input from a distant region region $S_j$ is delayed depending on the distance $d$ and conduction speed $v$. The global coupling scaling factor $G$ modulates the strength of these long-range connections.

3. **Local coupling**: Introduces surface geometry by describing how neighboring vertices k influence vertices l of region i.

$$L_i = \sum_{l \in i} \sum_{k}^{V} K(d_{kl}) \tag{4}$$

The term $L_i$ represents the short-range connections to region $i$ on the cortical surface. It sums up the influence of geometrically close regions. Here, $l$ denotes vertices within region $i$, and $k$ denotes vertices in the local neighborhood. The connectivity kernel $K(d_{kl})$ represents the coupling strength between vertices $k$ and $l$ based on their spatial distance $d_{kl}$.

1. **External Input** $I_{ext}$: Represents the direct input to the neural population in region $i$. This input can be an arbitrary temporal function, accounting for any external stimuli or perturbations of the neural population in that region.
2. **Noise** $\eta_i(t)$: Represents stochastic influences on the neural population in region $i$, reflecting random fluctuations and uncertainties in the neural **activity**. It can be additive but also multiplicative depending on the state $S_i$.

Incorporating all components, the state equation for region $i$ can be refined as:

$$\dot{S}_i = f_{\theta_i}(S_i(t-1)) + G\sum_{j=1}^{N} f_\lambda(S_j, S_i, g_{ij}) + \sum_{l\in i}\sum_{k}^{V} K(d_{kl})M_{k,i} + I_{ext} + \eta_i(t)$$

*1.1.3. Computational Knowledge*

We encode computational knowledge as a knowledge graph: a machine-readable network of typed entities and relations governed by a formal ontology and expressed with Web standards. This approach lets software interpret meaning (not just format), validate consistency, and answer semantic queries across models, data, and results. The main ingredients are:

1. **Ontology (schema)**: Defines classes, properties, and constraints that structure the graph. Computational ontologies instantiate domain concepts in a form computers can reason over, using RDF/OWL for semantics and SKOS for labels, synonyms, and acronyms. This complements rather than replaces traditional (philosophical) ontologies by providing an operational, machine-actionable vocabulary.
2. **Data (instances/facts)**: Concrete triples that populate the schema-for example, models, parameters (with units and ranges), equations, atlases, connectomes, experiments, and runs. These facts anchor identifiers to meanings and make entities queryable and comparable.
3. **Metadata (provenance and context)**: Who, what, when, and how for each entity and activity (authorship, timestamps, versions, software). Provenance follows the W3C PROV model so results remain traceable and reproducible across workflows.
4. **Inference and validation**: Lightweight reasoning and constraint checks derive implicit facts and guard consistency (e.g., subclass closure, unit compatibility, allowed value ranges). OWL/RDFS semantics and shape constraints provide automated checks without bespoke code.
5. **Querying and storage**: SPARQL exposes flexible retrieval and composition of entities (for example, "all excitatory parameters affecting gain in Jansen-Rit"). Scalable RDF stores and indices back responsive queries; APIs surface common queries to applications.
6. **Interoperability and integration**: Standards compliance (RDF, OWL, JSON-LD) and resolvable identifiers enable linking to external resources and round-tripping across systems via integration APIs-consistent with the broader Semantic Web.

In TVB-O, the ontology defines the primitives of brain network modeling, instances populate the graph from curated sources, PROV metadata records how results were produced, and SPARQL powers discovery and code export-keeping documentation, execution, and meaning in sync.

At web scale, knowledge graphs power widely used services (for example, Google's Knowledge Graph [3]) and are increasingly deployed in clinical decision support [4], underscoring their suitability for integrating scientific knowledge and workflows.

*1.2. SUPPLEMENTARY RESULTS*

*1.2.1. Mathematical Representation of Dynamical Systems in Ontology*

*1.2.2. Example Metadata Model*

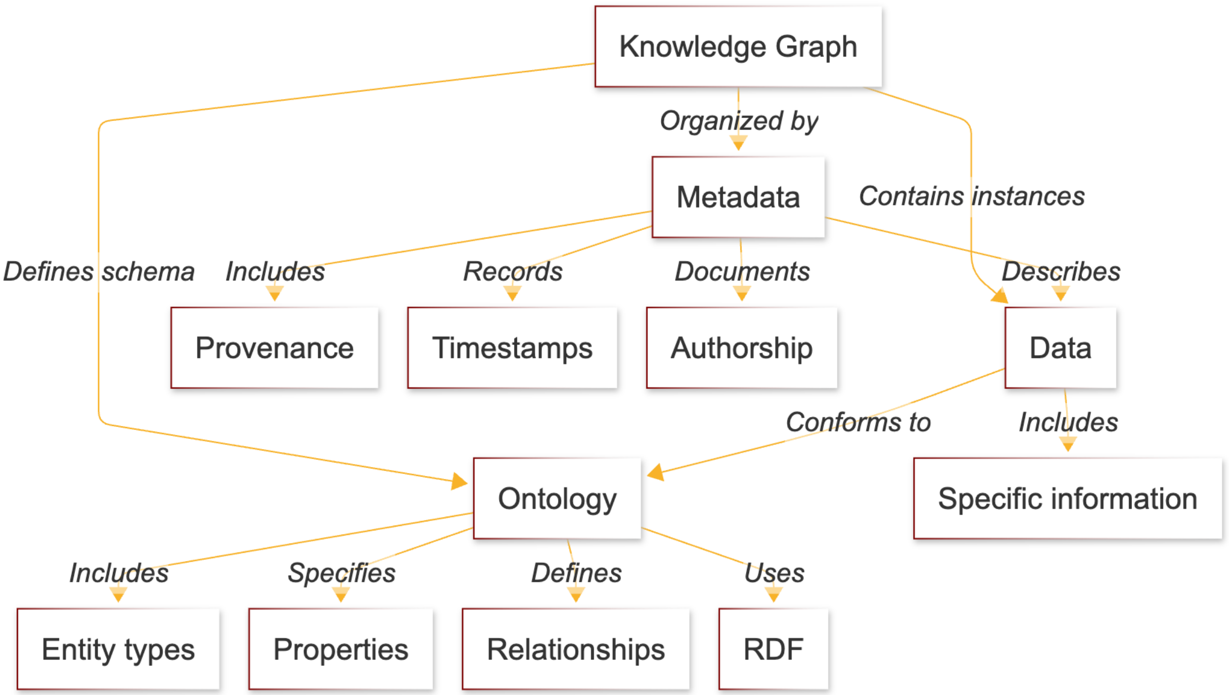

Figure 1: **Computational knowledge graph: components and flow.** The ontology defines the schema (classes, properties, constraints); instance data populate the graph with models, parameters, equations, atlases, connectomes, experiments, and runs; PROV metadata records who/what/how; lightweight reasoning and validation enforce consistency; SPARQL and APIs enable discovery and composition; and RDF/OWL/JSON-LD ensure interoperability. Together, these elements keep meaning, documentation, and execution aligned in TVB-O.

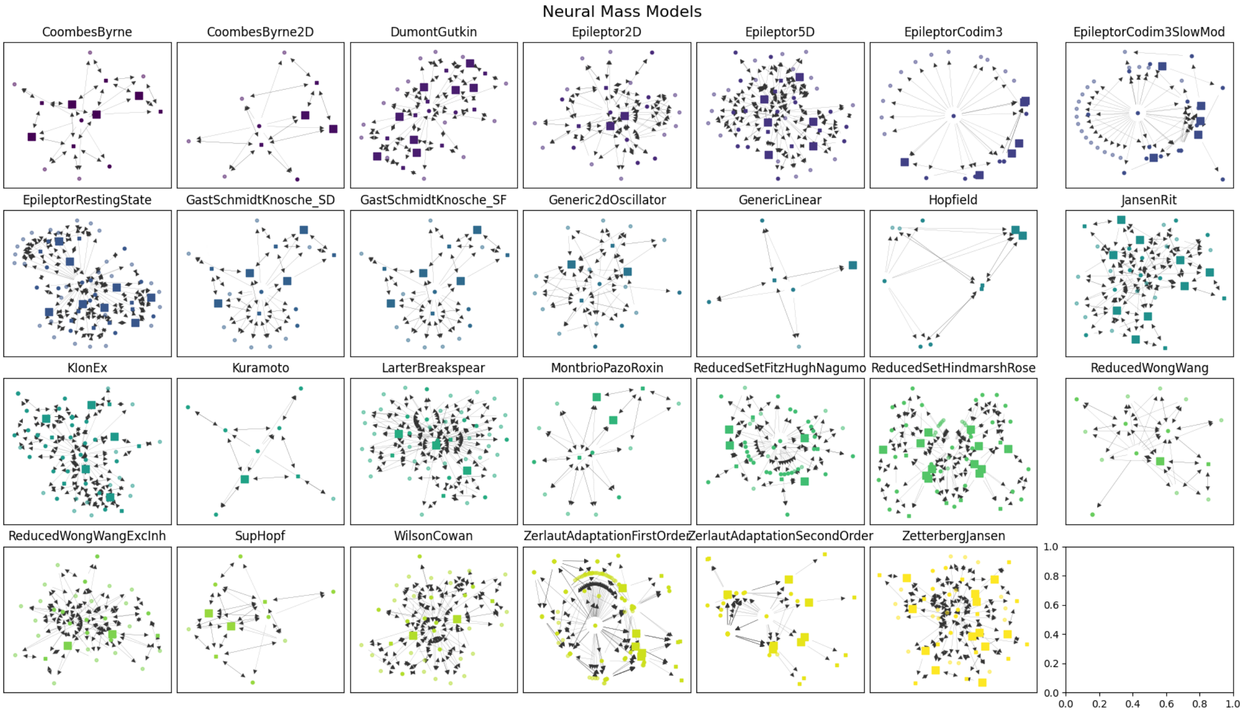

Figure 2: **Graphs of Neural Massmodels**. Mathematical relationships of Parameters and Variables.

```yaml
name: ModelJansen1995
has_reference: "Jansen1995"
label: "Jansen-Rit Neural Mass Model (single column)"
description: "JR model with canonical parameters; long-range input is injected via a
↪  coupling term. Output is pyramidal potential difference."
parameters:
A:
    value: 3.25
    unit: mV
B:
    value: 22
    unit: mV
C:
    value: 135
    explored_values:
    - 68
    - 128
    - 135
    - 270
    - 675
    - 1350
a:
    value: 0.1
    unit: ms^-1
b:
    value: 0.05
    unit: ms^-1
v0:
    value: 6
    unit: mV
e0:
    value: 0.0025
    unit: ms^-1
r:
    value: 0.56
    unit: mV^-1
p:
    value: 0.24
    unit: ms^-1
derived_parameters:
C1:
    equation:
    rhs: C
C2:
    equation:
    rhs: 0.8*C
C3:
    equation:
    rhs: 0.25*C
C4:
    equation:
```

```yaml
    rhs: 0.25*C
functions:
Sigm:
    definition: "JR sigmoid transformation"
    arguments:
    v:
        name: v
    equation:
    rhs: 2*e0 / (1 + exp(r*(v0 - v)))
coupling_terms:
c_glob:
    description: "Long-range input to the pyramidal population (pre-filter)."
state_variables:
y0:
    label: "Pyramidal membrane potential (position)"
    equation:
    rhs: y3
    unit: mV
y3:
    label: "Pyramidal EPSP drive (velocity)"
    equation:
    rhs: A*a*Sigm(y1 - y2) - 2*a*y3 - a**2*y0
    unit: mV*ms^-1
y1:
    label: "Excitatory interneuron membrane potential (position)"
    equation:
    rhs: y4
    unit: mV
y4:
    label: "Excitatory EPSP drive (velocity) with global input"
    equation:
    rhs: A*a*(p + C2*Sigm(C1*y0) + c_glob) - 2*a*y4 - a**2*y1
    unit: mV*ms^-1
y2:
    label: "Inhibitory interneuron membrane potential (position)"
    equation:
    rhs: y5
    unit: mV
y5:
    label: "Inhibitory IPSP drive (velocity)"
    equation:
    rhs: B*b*(C4*Sigm(C3*y0)) - 2*b*y5 - b**2*y2
    unit: mV*ms^-1
output_transforms:
- name: v_pyr
    description: "EEG-like output"
    equation:
    rhs: y1 - y2
    unit: mV
```

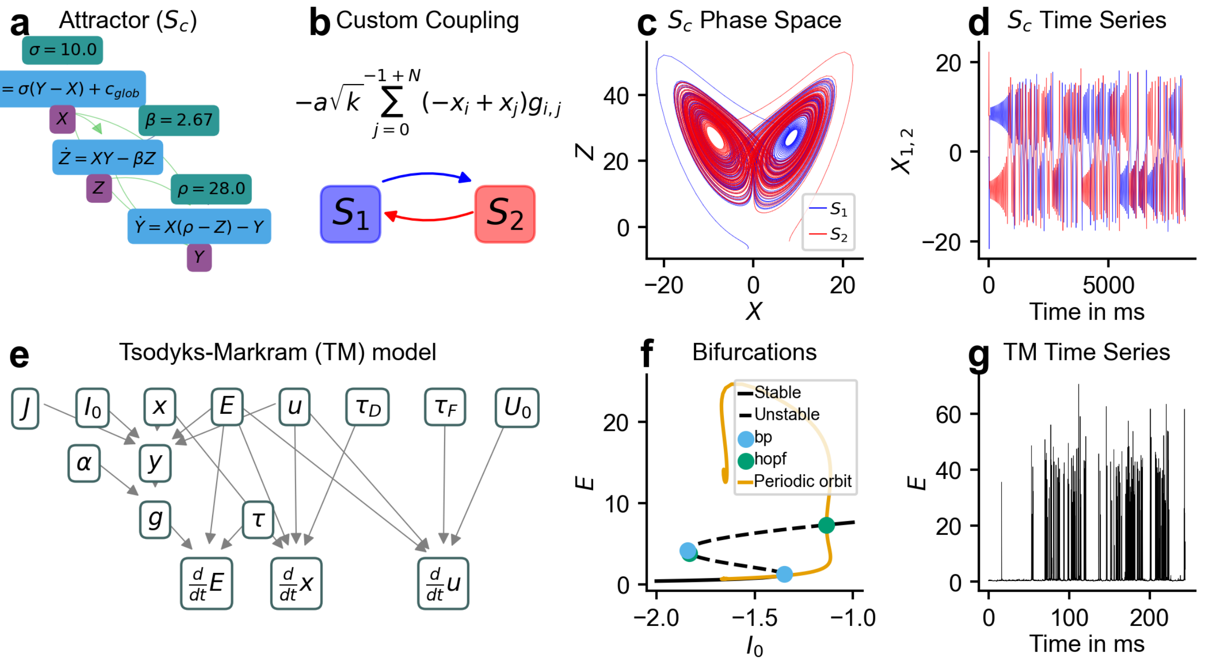

Figure 3: **Supplementary Figure S1 | Import custom models and coupling functions.** The first row (a-d) presents two coupled Lorenz attractors, their mathematical dynamics (a), a custom nonlinear coupling function (b), the classic phase-space visualization (c), and time series generated with TVB-O (d). The second row (e-g) shows the import of a Tsodyks-Markram model not yet in the ontology as a mathematical dependency graph (e) and a bifurcation analysis using Julia's BifurcationToolkit (f). Stable fixed points are solid black lines; unstable fixed points are dashed. Blue circles indicate saddle-node (bp) bifurcations, green circles indicate Hopf bifurcations, and orange curves correspond to periodic orbits. The simulated time series of the state variable E is shown in (g).

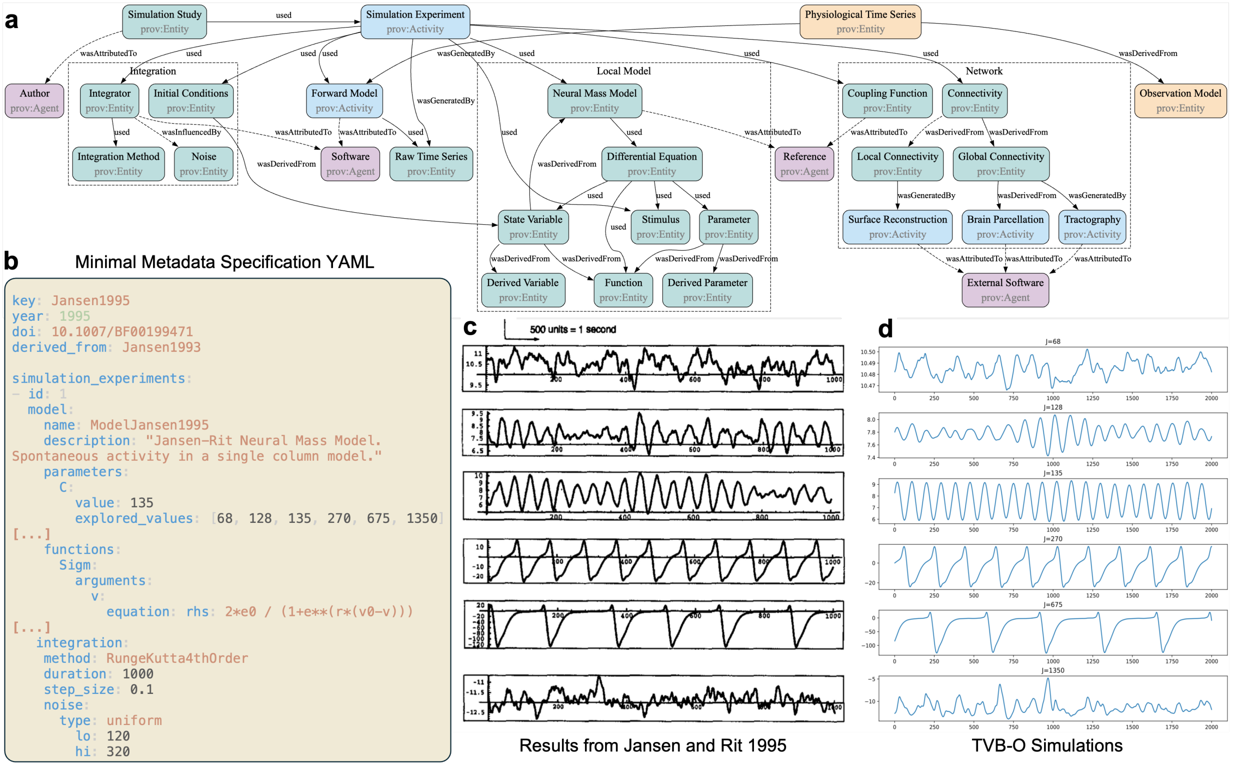

Figure 4: **Supplementary Figure S2 | Provenance graph and replication.** Panel (a) shows the PROV graph of a simulation with entities (green), agents (purple), and activities (blue) across the full dataflow. Panel (b) provides metadata excerpts for the Jansen-Rit single-column model. Panels (c-d) replicate the Jansen and Rit (1995) single-column model from a metadata file using model and integration settings; parameter C is swept from 68 (top) to 1350 (bottom), reproducing the dynamics described in the original publication.

*1.2.3. Example model import and custom coupling*

*1.2.4. Provenance model supports study replication*

The provenance model operationalizes end-to-end traceability by recording who, what, and how for each stage of a simulation, linking inputs, activities, and software to the resulting data. Applied to a classic Jansen-Rit study, these records allow the experiment to be re-instantiated directly from metadata, yielding trajectories and regime changes that agree with the original report across a sweep of the coupling parameter. Beyond a single example, the same pattern enables auditability-settings and versions are inspectable post hoc-and supports exact reruns or controlled variations without manual reconstruction of procedures.

*1.2.5. Supplementary Table S1. Unified list of neural population models and simulation study entries*

The unified table enumerates all ontology-aligned neural population models and curated simulation studies distributed with this release. It complements the narrative summary in Result 3 by providing provenance (TVB vs. external Julia source), dimensionality (state variable count), and availability of automated bifurcation templates.

Table 2: Unified list of Local Dynamics (Model) and SimulationStudy entries in TVB-O.

| NMM | UID | Parameters | Dimensions | Reference |
|---|---|---|---|---|
| CoombesByrne | TVBO:000126 | 5 | 4 | [5] |
| CoombesByrne2D | TVBO:000127 | 4 | 2 | [5] |
| DumontGutkin | TVBO:000170 | 14 | 8 | [6] |
| Epileptor2D | TVBO:000258 | 12 | 2 | [7, 8] |
| Epileptor5D | TVBO:000259 | 18 | 6 | [7, 9] |
| EpileptorRestingState | TVBO:000262 | 28 | 8 | [10, 9] |
| GastSchmidtKnosche_SD | TVBO:000521 | 9 | 4 | [11] |
| GastSchmidtKnosche_SF | TVBO:000522 | 9 | 4 | [11] |
| Generic2dOscillator | TVBO:000524 | 12 | 2 | [12, 13] |
| GenericLinear | TVBO:000525 | 1 | 1 | [14] |
| Hopfield | TVBO:000540 | 2 | 2 | [15, 16] |
| JansenRit | TVBO:000608 | 13 | 6 | [17, 18] |
| KIonEx | TVBO:000617 | 14 | 5 | [19, 20] |
| Kuramoto | TVBO:000645 | 1 | 1 | [21, 22, 23] |
| LarterBreakspear | TVBO:000650 | 32 | 3 | [24, 25, 26] |
| MontbrioPazoRoxin | TVBO:000677 | 7 | 2 | [27] |
| ReducedWongWang | TVBO:000765 | 8 | 1 | [28, 29] |
| ReducedWongWangExcInh | TVBO:000766 | 19 | 2 | [30] |
| SupHopf | TVBO:000814 | 2 | 2 | [31] |
| WilsonCowan | TVBO:000935 | 23 | 2 | [32, 33] |
| ZerlautAdaptationFirstOrder | TVBO:000942 | 58 | 5 | [34, 35] |
| ZetterbergJansen | TVBO:000944 | 18 | 12 | [36] |

*1.2.6. Supplementary Table S2. Full database inventory (all YAML records)*

The complete inventory of all curated YAML records across categories. This table is generated automatically from the pre-rendered CSV `db_inventory.csv` and reflects the current repository state at render time.

The full CSV is available in the project output at: `_output/extraction_results/db_inventory.csv`.

*1.2.7. Supplementary Table S3. TVB-O <-> Gene Ontology links*

All ontology cross-links between TVB-O classes and Gene Ontology terms used in the paper, with the predicate indicating the relation type. Counts reported in the main text are derived from this table during pre-render.

The full CSV is available in the project output at: `_output/extraction_results/go_links.csv`.

*1.2.8. Automated bifurcation analysis for exemplar models*

Bifurcation analysis in TVB-O is configured by a small set of controls and yields publication-ready diagrams with aligned time-series panels. The result is a reproducible process: continuations and time-domain sampling are derived from the same symbolic model specification, with equilibrium stability and periodic orbits tracked automatically. Two exemplars demonstrate the process: for the Generic 2-D Oscillator [37], a continuation over $a \in [-10, 20]$ with VOI $V$ identifies Hopf points and a family of periodic orbits, with representative trajectories illustrating quiescent, oscillatory, and steady regimes (see Figure 5 a); for Jansen–Rit [18], varying $\mu \in [-0.15, 0.5]$ exposes saddle-node and Hopf bifurcations and emergent periodic orbits around intermediate input, with trajectories spanning near-silent, damped, and oscillatory responses (see Figure 5 c).

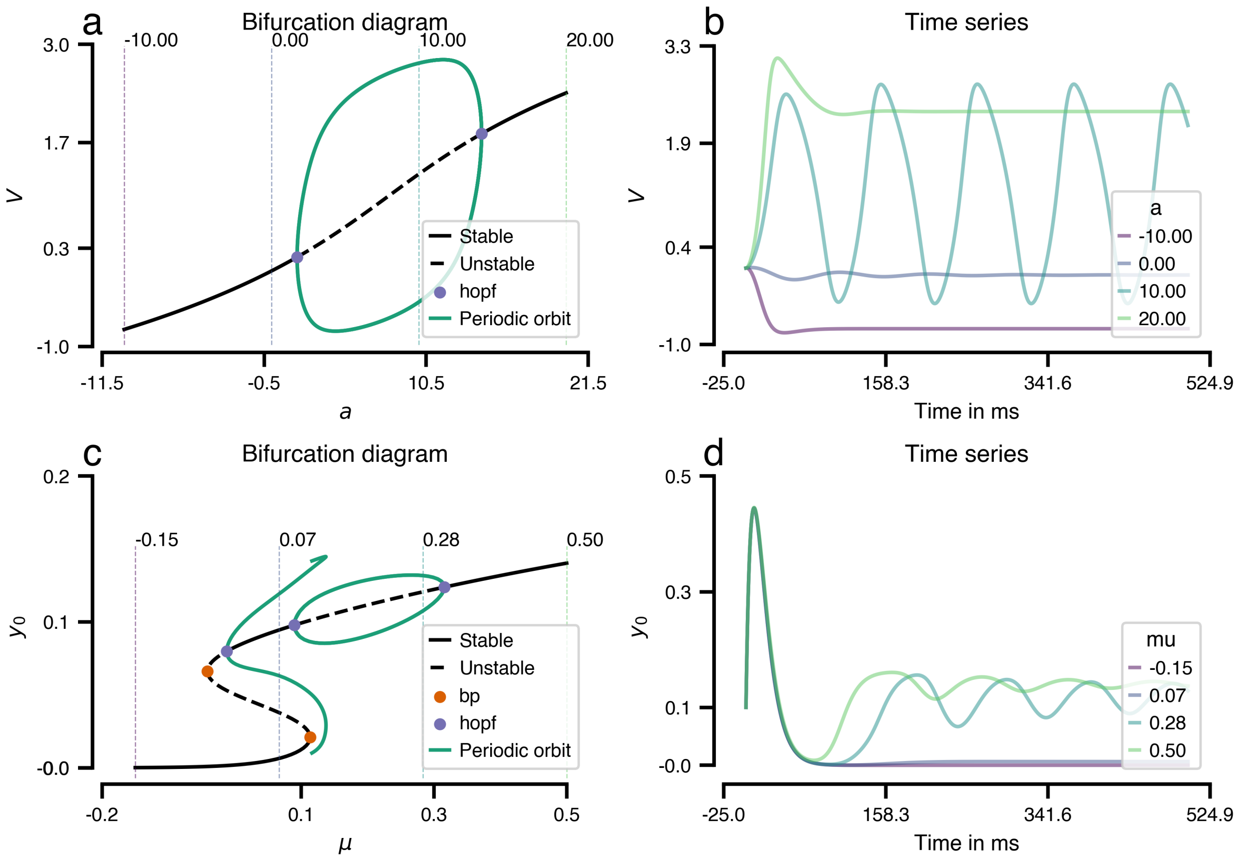

Figure 5: **Automated bifurcation analysis and time-series sampling.** Each row shows a one-parameter continuation and representative time series for: (top) a Generic 2-D Oscillator with control parameter $a$ and variable of interest $V$; (bottom) the Jansen-Rit model with control parameter $\mu$ and VOI $y_0$. Solid curves denote stable equilibria; dashed curves denote unstable equilibria. Periodic orbits, when present, are plotted by their minimum/maximum envelopes over the cycle. Filled markers indicate bifurcation points returned by the continuation backend: "bp" denotes saddle-node (fold) points on the equilibrium branch; "Hopf" denotes Andronov-Hopf bifurcations. Vertical dashed lines mark the parameter values used for the accompanying trajectories (four runs per model); the legend of the right panels lists the corresponding parameter values. Abbreviations: ICS, input control (continuation) parameter; VOI, variable of interest. Parameter ranges: $a \in [-10, 20]$ (top) and $\mu \in [-0.15, 0.5]$ (bottom). Continuations and periodic orbits were generated automatically from the symbolic model specification and cached; the plotting wrappers render both the bifurcation diagrams and the time-series panels from the same configuration.

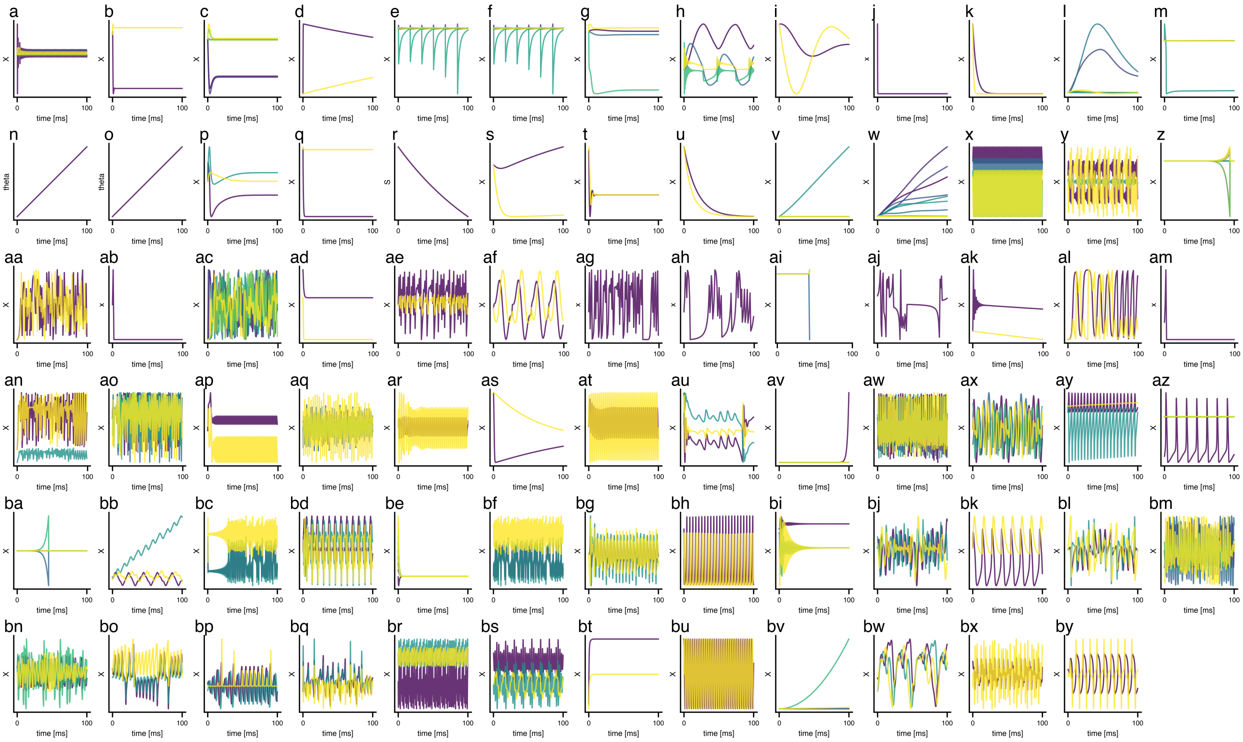

Figure 6: **Model library: repertoire of local dynamics.** Tiled gallery of all ontology-aligned local dynamics distributed with this release, generated directly by the code block above. Each small panel corresponds to one model; colored traces show its state variables (one color per state; colors have no semantic meaning). Simulations are single-node, uncoupled, and noise-free, using curated default parameters and zero initial conditions. Continuous-time models are integrated for 100 ms with a fixed step $\Delta t = 1$ ms; the first 5 ms are discarded as transient before plotting. Discrete-time models are iterated for 100 steps with $\Delta t = 1$ and plotted on the same 0-100 ms time base for comparability. Axes, limits, and aspect are standardized across panels, and a per-axes color cycle ensures distinct state traces. Panel letters identify models in the order discovered under database/models. The figure (~=3900 x 1800 px; 78 axes in this instance) is fully reproducible from the shown code, which scans database/models, loads each YAML with Dynamics.from_file, runs the simulation, and renders the traces.

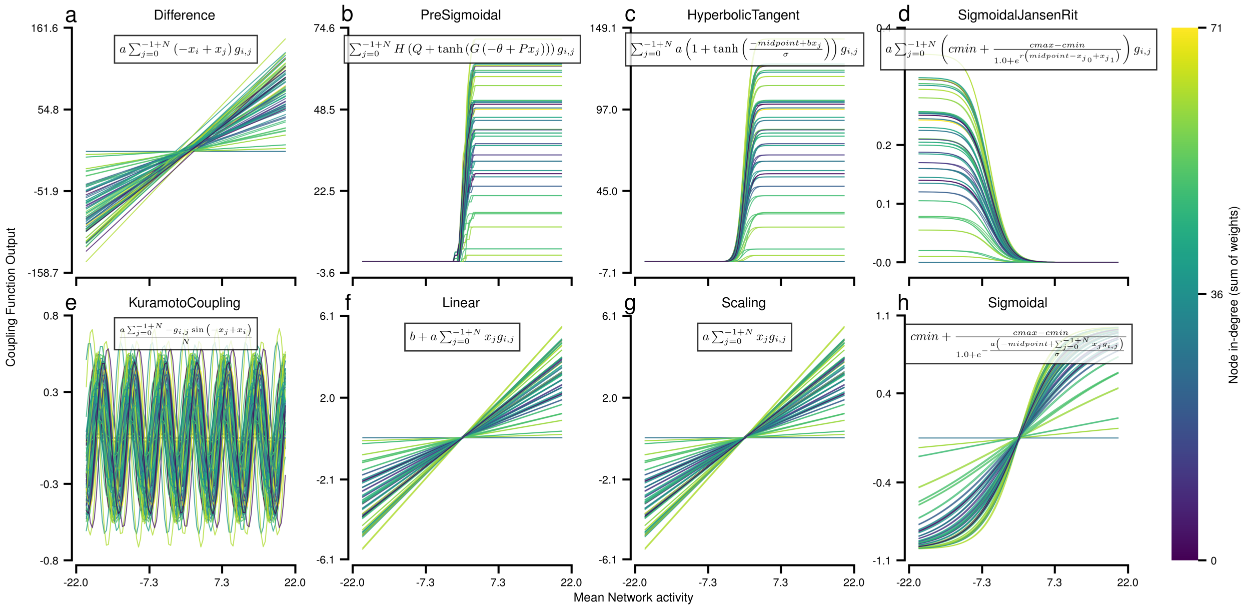

Figure 7: **Catalogue of coupling transforms and operating regimes.** Each panel (a-h) shows the input-output relation of a built-in coupling function used to aggregate afferent activity across a network: Difference, Pre-Sigmoidal, Hyperbolic-Tangent, Sigmoidal-Jansen-Rit, Kuramoto phase coupling, Linear, Scaling, and Sigmoidal. Curves correspond to individual nodes; color encodes node in-degree (sum of incoming weights) to illustrate how typical operating ranges vary with network context. Insets display the analytical forms rendered from the symbolic specification. Together, these transforms span monotone linear, saturating, and oscillatory regimes and can be interchanged in experiments with explicit pre-/post-synaptic semantics.

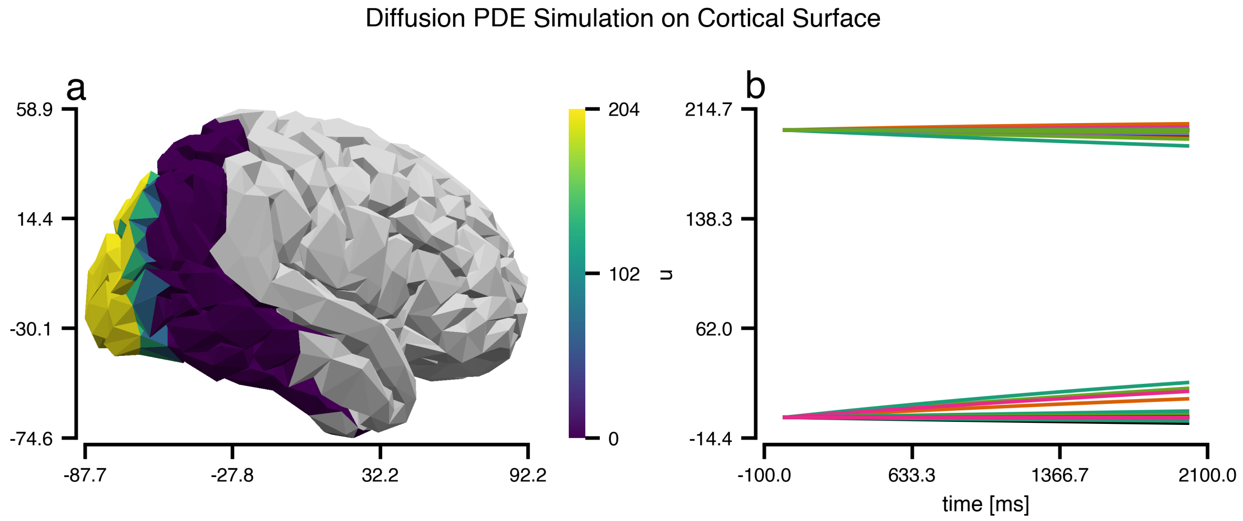

Figure 8: **Surface diffusion PDE driven from a TVB-O specification.** A minimal diffusion equation on the cortical surface is executed from an ontology-aligned experiment description. Panel (a) shows the scalar field on a right-hemisphere mesh at the final time; the initial condition is localized to a temporal patch and diffuses over the triangulated surface. Panel (b) displays representative vertex-wise trajectories. The solver is generated from the same specification and uses first-order finite elements on triangles with an implicit-Euler time step; GIFTI meshes are loaded directly and Dirichlet boundaries applied on the outer surface. This demonstration underscores that PDEs can be configured, executed, and reproduced from the same machine-readable description that drives network simulations.

###### *1.2.9. Model Library - Repertoire of dynamical systems*

###### *1.2.10. Library of coupling functions*

###### *1.2.11. Surface-based PDE simulation*

###### *1.2.12. Annotated database of normative structural connectivity*

Table 3: Atlas repertoire used to derive normative structural connectivity matrices (weights and tract lengths). Region counts reflect the distributed versions in TVB-O; Schaefer provides multi-resolution sets (we use the 17-network and 1000-parcel resolutions). All atlases are supplied with versioned metadata and region centers enabling cross-atlas mapping and delay derivation.

| Atlas | Regions (n) | Parcellation Type | Reference |
|---|---|---|---|
| Yeo networks | 17 | Functional intrinsic connectivity networks | [38] |
| Desikan–Killiany | 68 | Gyral anatomical segmentation | [39] |
| Destrieux | 148 | Sulco-gyral anatomical segmentation | [40] |
| HCP-MMP1 | 360 | Multi-modal (architecture, function, connectivity, myelin) | [41] |
| Schaefer | 1000 | Local–global functional parcels | [42] |

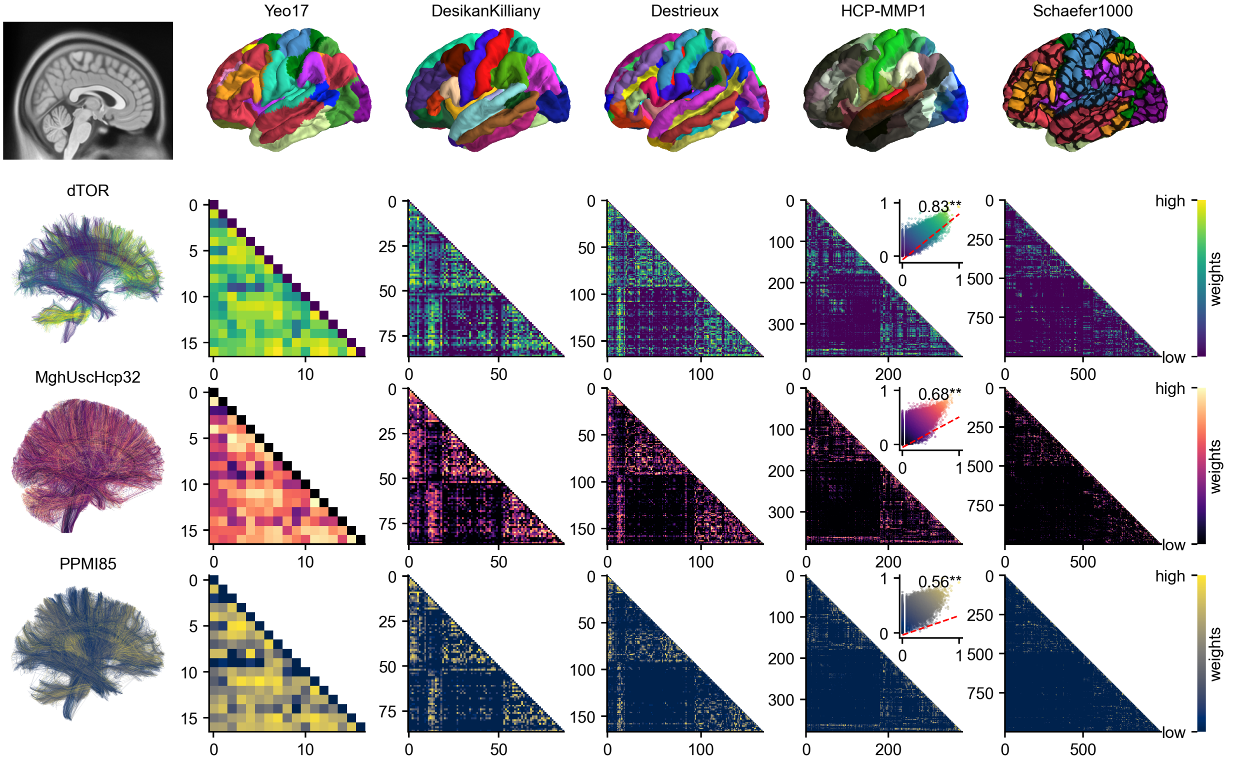

Figure 9: **Normative structural connectivity.** Overview of normative structural networks provided by TVB-O. Rows show connectivity matrices derived from multiple normative tractograms parcellated by standard atlases with increasing granularity; scatterplots for HCP-MMP1 summarize weight and length distributions.

TVB-O distributes a curated, ready-to-use library of normative structural brain networks derived from five standard parcellation atlases (Table 3) and three underlying normative tractograms — two from healthy cohorts (dTOR, MghUscHCP32) [43, 44, 45] and one with Parkinson's Disease (PPMI) [46, 47]. For each atlas–tractogram combination, TVB-O provides the region-wise connectivity matrices with symmetric weights (Figure 9) and mean tract lengths with explicit units, enabling delays to be derived via a physiologically plausible conduction speed. These normative matrices follow a BIDS-like organization, carry minimal provenance, and are validated for internal consistency (labels, shapes, and units). We use them throughout the paper, for example, the Yeo-17 example shown in Figure 7 of the main manuscript. Full curation, unit conventions, and validation procedures are detailed in Methods (Section 1.3.5).

##### *1.2.13. Benchmarking with state-of-the-art simulators (TVB)*

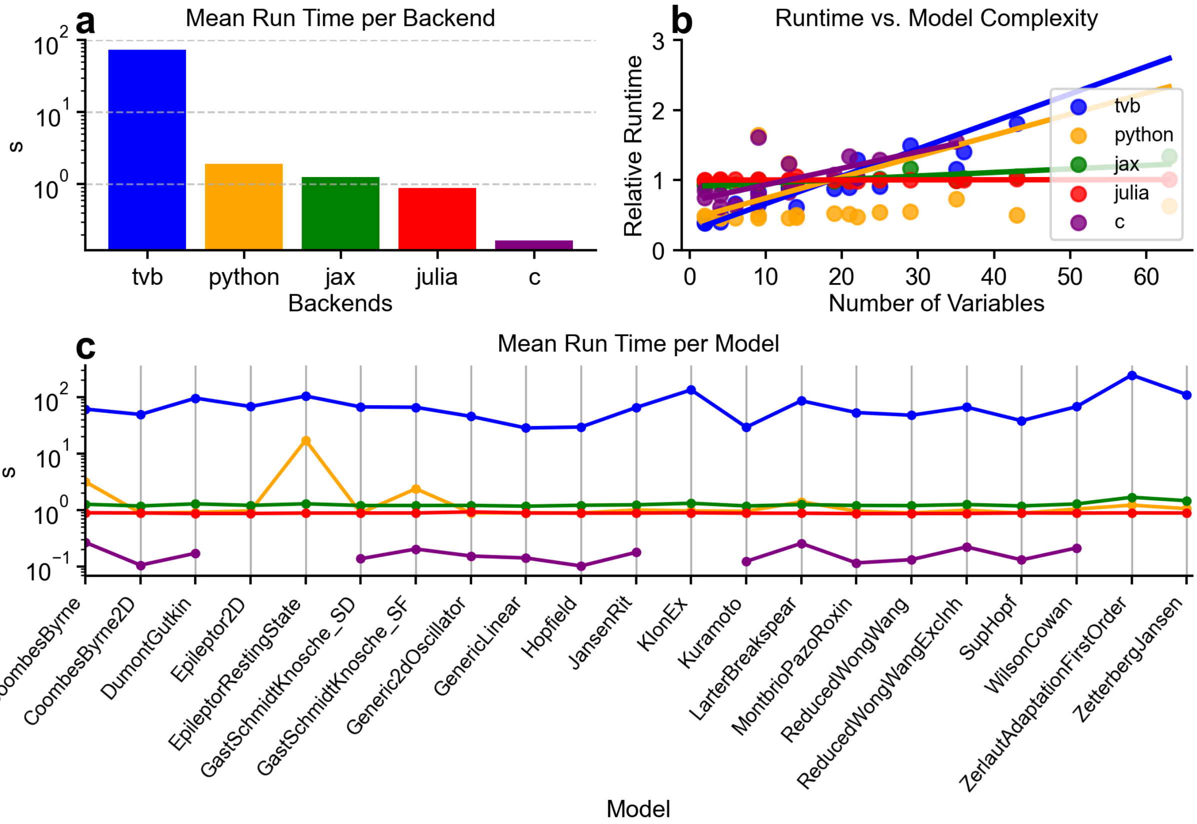

Figure 10: **Runtime comparison across backends (10-s simulated, $\Delta t = 1$ ms).** (a) Mean wall-clock time per backend, aggregated over models (log scale). (b) Relative runtime versus model complexity (number of variables + parameters) with per-backend regression lines. (c) Mean runtime per model across backends (log scale).

Under identical numerical settings, the C baseline is consistently fastest across models, followed by Julia and JAX without JIT, then the pure-Python reference; the TVB path is slowest on average (panel a). Across models, runtimes vary over orders of magnitude on a log scale, but JAX and Julia show weak dependence on the simple complexity index, whereas Python exhibits a mild positive trend and TVB a steeper slope (panel b), reflecting higher overheads as model size increases. Per-model profiles are stable across the library, with the ordering of backends largely preserved (panel c). Configurations that failed to execute were skipped, and high-dimensional variants not broadly supported (e.g., Epileptor5D) were excluded from aggregates. Timings reflect full per-call execution in each pathway and exclude I/O and plotting; identical equations and settings were used across backends to ensure workload parity (see Methods S2.4).

##### *1.2.14. Integration of the AtOM ontology for brain atlas metadata*

We adopt a standardized atlas metadata profile to make parcellations and their coordinate frames first-class, machine-readable components of simulation entries. In practice, this captures four complementary elements: a citable atlas record with versioning and a link to its reference coordinate space; a description of that coordinate space, including units, orientation, and origin; a controlled parcellation terminology that enumerates regions with stable labels, names, hierarchy, hemisphere, representative center coordinates, colors, and cross-ontology identifiers; and region entities as the addressable units used by models and stimuli. When a structural network is selected, its parcellation anchors both an atlas version (terminology and reference metadata) and a coordinate space (units and axes), enabling unambiguous translation between region identifiers and matrix indices and ensuring that delays derived from tract lengths and conduction velocity are interpreted in the declared spatial units. For FAIR reuse, we validate a minimal set of fields at ingest-atlas name and version (with a stable digital identifier), coordinate-space units and axes orientation, a complete list of parcellation entities with stable labels/names and external identifiers, and region center coordinates-and export them alongside experiments. This alignment supports reproducible network assembly, precise cross-atlas mappings, and round-trips to EBRAINS/Siibra without exposing users to implementation details.

##### *1.3. SUPPLEMENTARY METHODS*

Flowchart of the curation pipeline: from more than one million GO concepts to a screened set of 215 electrophysiology-relevant processes linked across TVB-O layers (biological processes -> BNM components -> models/parameters), summarizing filtering, expert review, and ontology-linking criteria.

###### *1.3.1. Automated bifurcation analysis pipeline*

We provide an automated pipeline for codimension-1 continuation that operates directly on the model's symbolic specification. Given a system f(x, p) = 0 with state x and control parameter p (ICS), TVB-O derives a continuation program from the same equations and parameter values used for simulation and executes it to obtain equilibrium branches x⋆(p) over a user-defined interval. Linear stability is assessed along each branch from the Jacobian spectrum and, when enabled, families of periodic orbits are computed and recorded. All results are serialized to machine-readable files.

The backend is Julia's BifurcationToolkit, invoked via a self-contained program emitted from the symbolic form of the model. Using a single source for simulation, reporting, and continuation eliminates documentation-implementation drift and keeps parameter names and units consistent. Artifacts (equilibria, stability, limit-cycle summaries) are cached under deterministic file names keyed by the model identifier and continuation settings, ensuring that repeated analyses with identical inputs reproduce the same outputs and that cached results are reused where available.

For interpretability, the pipeline augments each diagram with short trajectories at a small set of marked parameter values spanning qualitatively distinct regimes. These time-domain traces link changes in stability to observable dynamics of a chosen variable of interest (VOI).

In this work we considered two exemplars. For a Generic 2-D Oscillator we varied the parameter $a$ on [-10, 20] and plotted $V$ as VOI; for the Jansen-Rit model we varied the background input $\mu$ on [-0.15, 0.5] and plotted the pyramidal output $y_0$. Diagrams display stable (solid) and unstable (dashed) equilibria and, where present, periodic orbits; vertical markers indicate the parameter values used for the accompanying trajectories (Supplementary Figure 5).

All panels of Supplementary Figure 5 can be regenerated with the provided script (`manuscript/code/`). Because artifacts are deterministically named from the model and settings, re-execution on another system yields identical file names and figures when using the same backend versions.

###### *1.3.2. Supplementary Method: Generating the coupling-function library (Supplementary Figure 7)*

Scope and definitions. We assembled a library of eight commonly used coupling transforms-Difference, Pre-Sigmoidal, Hyperbolic-Tangent, Sigmoidal-Jansen-Rit, Kuramoto phase coupling, Linear, Scaling, and Sigmoidal-captured as ontology-aligned records within TVB-O. Each record specifies a closed-form expression and a default parameterization together with a short description of its intended use.

Evaluation. To characterize these transforms independently of any particular study, each expression was evaluated across a standardized input range representative of large-scale brain models (x in [-22, 22]). Curves in the figure depict node-wise responses when the transform is applied to afferent activity under default semantics (pre-synaptic transform, weighted summation across inputs, optional post-synaptic transform). Color encodes node in-degree (sum of incoming weights) to indicate typical operating regimes in realistic networks. Kuramoto coupling is shown in its canonical sinusoidal form based on phase differences and averaged over afferents.

Presentation and comparability. Insets display the analytical form of each transform. Axes and limits are shared across panels to support visual comparison between monotone linear, saturating, and oscillatory behaviours. Parameter values are fixed at their curated defaults for all nodes; no per-node fitting was performed.

Reproducibility. Because the definitions and defaults are versioned within the resource and linked to stable identifiers, the panel shapes and legends can be regenerated from the same records without reliance on bespoke code.

Coupling semantics and exemplary forms. Throughout TVB-O, inter-regional input to node i is expressed as a pre/post transform around a weighted sum of afferents

$$\begin{aligned}\psi_j &= f_{\text{pre}}(x_j),\\ s_i &= \sum_{j=1}^{N} g_{ij}\,\psi_j,\\ c_i &= f_{\text{post}}(a\,s_i + b),\end{aligned}$$

where $x_j$ is the presynaptic signal at node $j$, $g_{ij}$ is the connection weight, and $a, b$ are gain and offset. The library instantiates this template with common choices; typical examples include:

- Difference (anti-symmetric drive):
$$c_i = a\sum_j g_{ij}\,\big(x_j - x_i\big).$$
- Linear:
$$c_i = a\sum_j g_{ij}\,x_j + b.$$
- Scaling (gain-only linear):
$$c_i = a\sum_j g_{ij}\,x_j.$$
- Hyperbolic-tangent (smooth saturation):
$$c_i = a\left[1 + \tanh\!\left(\tfrac{s_i - m}{\sigma}\right)\right],$$
with midpoint $m$ and scale $\sigma$.
- Sigmoidal (logistic, bounded between $c_{\text{min}}$ and $c_{\text{max}}$):
$$c_i = c_{\text{min}} + \frac{c_{\text{max}} - c_{\text{min}}}{1 + e^{-\frac{s_i - m}{\sigma}}}.$$
- Sigmoidal-Jansen-Rit (population-level logistic):
$$c_i = a\,\frac{2e_0}{1 + e^{r\,(v_0 - s_i)}},$$
with slope $r$, midpoint $v_0$, and scale $2e_0$.

- Pre-sigmoidal (nonlinearity before summation; example):

$$\psi_j = \tanh\big(G\,(x_j - \theta)\big), \quad c_i = \sum_j g_{ij}\,\psi_j.$$

- Kuramoto phase coupling:

$$c_i = \frac{a}{N}\sum_{j=1}^{N} g_{ij}\,\sin\big(\phi_j - \phi_i\big).$$

Parameter symbols follow the curated defaults linked to each definition.

*1.3.3. Supplementary Method: Surface diffusion PDE experiment and rendering (Supplementary Figure 8)*

Experiment design. The diffusion demonstration is expressed as an ontology-aligned experiment that specifies: a cortical surface domain represented by a triangular mesh (GIFTI format), a single scalar state variable u, homogeneous (zero) Dirichlet boundary conditions on the outer surface, and a diffusion operator with coefficient $D = 1.0x10^{-6}$. The simulation uses a fixed time step of $\Delta t = 0.5$ and runs for 100 ms (200 steps).

Numerical approach. A finite-element scheme on linear triangular elements advances the heat equation $u_t = D \cdot \Delta u$ using a backward (implicit) Euler integrator. Essential boundary values are enforced on the outer surface. This choice yields a stable update for the chosen step size and provides a natural discretization on irregular cortical meshes commonly used in neuroimaging.

Initialization and stimulus. To elicit a non-trivial evolution under homogeneous boundaries, the initial condition is spatially localized: vertices with y < -60 mm are set to 200 (arbitrary units), elsewhere 0. No external forcing is applied; the observed spread is therefore driven solely by diffusion on the surface geometry.

Visualization. The left panel maps the final field to the right-hemisphere surface; the right panel shows representative vertex-wise trajectories. For interpretability across time, a single color scale is applied using global minimum/maximum values computed over the full time series. The mesh used for visualization is the same as for the computation, ensuring geometric consistency.

Reproducibility. The entire workflow-domain, operator, numerical settings, and initialization-is captured in a single, machine-readable experiment description. Because meshes are read from standard surface files and the solver is generated directly from the specification, the figure can be reproduced without manual re-implementation.

Governing equations and discretization. The continuous problem is

$$\begin{aligned} \partial_t u(t,\mathbf{x}) &= D\,\Delta u(t,\mathbf{x}) && \text{in } \Omega, \\ u(t,\mathbf{x}) &= 0 && \text{on } \partial\Omega, \\ u(0,\mathbf{x}) &= u_0(\mathbf{x}), \end{aligned}$$

with $\Omega$ the cortical surface and $\partial\Omega$ its boundary. The backward-Euler finite-element update reads: find $u^{n+1} \in V_0$ such that for all test functions $v \in V_0$,

$$\big(u^{n+1}, v\big) + \Delta t\, D\,\big(\nabla u^{n+1}, \nabla v\big) = \big(u^n, v\big) + \Delta t\,\big(f^n, v\big),$$

which leads to the linear system

$$\big(M + \Delta t\, D\, K\big)\,\mathbf{u}^{n+1} = M\,\mathbf{u}^n + \Delta t\, M\,\mathbf{f}^n,$$

where $M$ and $K$ are the mass and stiffness (Laplace) matrices and essential boundary values are applied on $V_0$.

*1.3.4. Models used in Use Case #2: Comparing Models*

This section summarizes the two neural-mass models compared in Use Case #2 (Figure 5 in the main manuscript). Content is adapted from auto-generated TVB-O model reports to match manuscript style; full specifications and PDF reports are provided in the repository (Methods 3.4.3-3.4.4).

*1.3.4.1. Jansen-Rit neural mass model (YAML ID: Jansen1995).*

Three interacting populations (pyramidal, excitatory interneuron, inhibitory interneuron) are coupled via second-order synaptic kernels; a sigmoid maps membrane potential to firing rate [18].

Derived parameters

$$C_1 = C, \quad C_2 = 0.8\,C, \quad C_3 = 0.25\,C, \quad C_4 = 0.24\,C.$$

Transfer function

$$\mathrm{Sigm}(v) = \frac{2e_0}{1 + e^{r\,(v_0 - v)}}.$$

State equations

$$\begin{aligned}
\dot{y}_0 &= y_3, & \dot{y}_3 &= -a^2 y_0 - 2ay_3 + Aa\,\mathrm{Sigm}(y_1 - y_2),\\
\dot{y}_1 &= y_4, & \dot{y}_4 &= -a^2 y_1 - 2ay_4 + Aa\,\big(p + C_2\,\mathrm{Sigm}(C_1 y_0)\big),\\
\dot{y}_2 &= y_5, & \dot{y}_5 &= -b^2 y_2 - 2by_5 + BC_4 b\,\mathrm{Sigm}(C_3 y_0).
\end{aligned}$$

Parameters

| Parameter | Value | Unit |
|---|---|---|
| $C$ | 135.0 | - |
| $A$ | 3.25 | mV |
| $B$ | 22.0 | mV |
| $a$ | 0.10 | ms^-1 |
| $b$ | 0.05 | ms^-1 |
| $v_0$ | 6.0 | mV |
| $e_0$ | 0.0025 | ms^-1 |
| $r$ | 0.56 | mV^-1 |
| $p$ | 0.24 | - |

References: [17, 18].

*1.3.4.2. Reduced Wong-Wang excitatory-inhibitory model (YAML ID: ReducedWongWangExcInh).*

Two state variables represent excitatory and inhibitory synaptic gating $(S_E, S_I)$. Inputs combine local recurrence, inter-regional coupling, and background drive; non-linear response functions map inputs to activity [30].

Selected derived quantities

$$
\begin{aligned}
J_{NS_e} &= J_N S_E, \qquad \text{coupling} = G J_N \left(c_{\text{glob}} + S_E c_{\text{local}}\right),\\
x_E &= -b_E + a_E\big(I_{ext} + \text{coupling} + I_0 W_E + J_{NS_e} w_p - J_I S_I\big),\\
x_I &= -b_I + a_I\big(J_{NS_e} - S_I + I_0 W_I + \text{coupling}\,\lambda\big),
\end{aligned}
$$

$$
H_E = \frac{x_E}{1 - e^{-d_E x_E}}, \quad H_I = \frac{x_I}{1 - e^{-d_I x_I}}.
$$

State equations

$$
\begin{aligned}
\dot{S}_E &= -\tfrac{S_E}{\tau_E} + H_E\,\gamma_E\,(1 - S_E),\\
\dot{S}_I &= H_I\,\gamma_I - \tfrac{S_I}{\tau_I}.
\end{aligned}
$$

Parameters

| Parameter | Value | Unit | Note |
|---|---|---|---|
| $G$ | 2.0 | - | Global coupling |
| $I_{ext}$ | 0.0 | - | External drive to $E$ |
| $I_0$ | 0.382 | nA | Background input |
| $J_N$ | 0.15 | - | Excitatory gain |
| $J_I$ | 1.0 | - | Inhibitory feedback to $E$ |
| $W_E$ | 1.0 | - | Input weight to $E$ |
| $W_I$ | 0.7 | - | Input weight to $I$ |
| $a_E$ | 310.0 | nC^-1 | Response slope ($E$) |
| $a_I$ | 615.0 | nC^-1 | Response slope ($I$) |
| $b_E$ | 125.0 | Hz | Response shift ($E$) |
| $b_I$ | 177.0 | Hz | Response shift ($I$) |
| $d_E$ | 0.16 | s | Response scale ($E$) |
| $d_I$ | 0.087 | s | Response scale ($I$) |
| $\gamma_E$ | 6.41e-4 | - | Kinetics ($E$) |
| $\gamma_I$ | 0.001 | - | Kinetics ($I$) |
| $\lambda$ | 0.0 | - | Inhibitory coupling scale |
| $\tau_E$ | 100.0 | ms | NMDA decay |
| $\tau_I$ | 10.0 | ms | GABA decay |
| $w_p$ | 1.4 | - | Recurrent weight ($E$) |

Reference: [30].

*1.3.5. METHOD for result S2.1: Creation of the annotated connectome database (supplementary)*

We compiled a compact, BIDS-inspired library of normative structural connectivity to support reproducible network simulations and figure generation. The resource pairs versioned voxelwise parcellations (with atlas/space metadata and stable region labels) with region-wise connectivity matrices of weights (streamline counts) and mean tract lengths (millimeters). Where region centers are unavailable, representative coordinates are estimated with standard neuroimaging procedures; when helpful, we also provide a consecutively indexed gray-matter-focused variant that preserves traceability to original labels. Matrices are derived from tractography parcellation and are symmetric with zero diagonals by construction. Delays used in simulations are obtained as lengths divided by a conduction speed, with a default of 3.0 mm/ms to yield delays in milliseconds consistent with atlas units. Each dataset is indexed by atlas and a descriptor indicating the tractography pipeline or cohort. We validate internal consistency between segmentations, metadata, and matrix shapes, flagging any discrepancies without silent relabeling. Minimal provenance is recorded, and deterministic visualization settings ensure repeatable figures. Units follow neuroimaging conventions (centers in native atlas coordinates, lengths in millimeters, speed in mm/ms, delays in ms). This resource underlies Supplementary Fig. S3 and the Yeo-17 example shown in Figure 7 of the main manuscript.

###### *1.3.6. METHOD for result S2.4: Benchmarking and validation with The Virtual Brain*

We assessed execution speed across supported computational pathways by measuring wall-clock time to integrate each curated neural-mass model for a fixed simulated duration of 10 seconds at a step size of 1 ms under identical numerical settings. Five backends were exercised: a TVB-native path, a pure-Python reference, a JAX implementation evaluated without just-in-time compilation to avoid warm-up effects, a Julia-based path, and a C baseline where available. For every model-backend combination, a single run was executed and its runtime recorded; configurations that failed to execute were skipped, and models lacking complete backend coverage (for example, high-dimensional variants not supported on all paths) were excluded from aggregate summaries. To balance reproducibility with efficiency, timings were memoized per model, backend, and duration and reused on subsequent renders (the C measurement was recomputed on each run to avoid potential staleness). Results are summarized as (i) the mean runtime per backend across models and (ii) the mean runtime per model across backends, both displayed on a logarithmic scale. To relate performance to problem size, we regressed within-backend relative runtime against a simple complexity index combining the number of state variables and parameters for each model. Timings reflect the full per-call execution in each pathway (including any setup or compilation that occurs at invocation) and exclude I/O and plotting. Identical equations and numerical settings were used across backends to ensure workload parity.
